# Fragment size and diversity of mulches affect their decomposition, nutrient dynamics, and soil microbiology

**DOI:** 10.1101/2022.07.21.500764

**Authors:** Dimitrios Gaitanis, Martin Lukac, Mark Tibbett

## Abstract

Plant-based mulch has been proposed as a sustainable way of maintaining soil fertility. However, the role of mulch diversity, quality, and size in decomposition dynamics, and their effect on crop yield are not fully explored. We investigated how mulch quality, proxied by the constituent plant species diversity, and residue size drive mulch decomposition, crop nutrition, and yield.

A rhizotron experiment was set up with barley as a model crop, with the addition of mulch of two particle sizes (1.5 and 30 cm) and four different plant residue mixes (17, 12, 6, and 1 species) in a fully factorial design. Soil nutrient dynamics were measured at advanced decomposition stages, together with residue quality, arbuscular mycorrhizal fungal (AMF) root colonization, and crop yield.

Residue mass loss was significantly affected by its chemical composition. *Long residues* retained significantly higher C and N content, than *short residues*. Crop yield was not affected by residue type or size. Residue size significantly affected barley growth rate, influencing seed protein content. Soil available K was significantly increased by residues with a higher initial C:N ratio. *Short residues* resulted in higher soil Zn. Residues of higher diversity resulted in higher AMF root colonization of the barley plants.

Generally, l*ong residue* mulches maintain soil fertility for a longer period than *short* ones, without a deleterious effect on crop yield. Further investigation should evaluate the effect of continuous application of *long residue* mulches on soil fertility and microbial populations.

## Introduction

Plant-based mulch can significantly affect the physical, chemical, and biological properties of agricultural soils [1–4]. Mulches can provide physical protection from soil erosion and enhance and maintain soil fertility, particularly in conservation tillage, and organic farming systems [5–7]. Sustainability in agroecosystems is typically defined by the reliance on local resources, and diversity conservation [8]. Therefore, mulch from cover crops and crop residues can contribute to sustainability through the promotion of soil fertility and the diversity of soil-dwelling organisms [9]. However, in contrast to mineral fertilizers, the release of nitrogen from organic sources sometimes is unpredictable and much more susceptible to variations in environmental conditions [10]. The effect of mulching on soil carbon (C) and nitrogen (N) – as indicators of soil quality - is not well understood [11]. Generally, the greater the mass of residues, the higher the soil C content, yet the decomposition curve of N is not so straightforward and is affected by a number of soil conditions [12].

The timing of nutrient release relative to crop demand through the growing season has been found to be an important issue when using mulch. Su et al. [4] found increased N uptake and growth by plants during the early stages of their development, soon after mulching application. They note that these effects were diminishing with time, probably due to enhanced ammonia volatilization. However, soil N, mainly NH_4_^+^ and organic N availability, does not always increase immediately after mulch application. Siczek et al. [13] showed that N availability after mulch application could be limited during crop vegetative stages when soil nitrate content is low. Moreover, a rapid decomposition is not always beneficial, as it may provide nutrients too early, leading to a lack at latter stages [9].

Concentrations of available nutrients can also be dependent on the number of applications of plant mulch. Pavlu et al. [14] observed increasing concentrations of available P and K in ascending order from one to three annual applications. P concentrations are typically higher in topsoil due to the decomposition of organic materials and plant residues, to rapid fixation by soil particles, and to immobilization by microorganisms [15]. Moreover, higher seed protein content in soybean was reported with mulch application, whereas there were no measurable effects on yield in comparison to no mulch [13].

Residue diversity can stimulate soil ecosystem services but there is a paucity of knowledge on the type (plant traits) and the quantity (species richness) of the diversity required [16,17]. When plant residues are derived from a mixture of plant species, their decomposition process can be either faster (synergistic effect), slower (antagonistic effect), or proportional to constituents (additive), depending on the individual plants which participate [18]. Crop residues of a mixture of different plant species may increase plant residue-derived C assimilation by soil microbes in the early stage of decomposition, than residues of single plant species, due to functional complementarity which results in reduced competition between microbial communities [19]. Substrates of different chemical composition are decomposed by different groups of microbes, using different enzymes [20]. Therefore, residues with high plant diversity favour microbial diversity [21] and arbuscular mycorrhizal fungal (AMF) diversity [22].

The decomposition of plant residues can be affected by the amount of contact between soil particles and plant material and consequently by the size of plant residue per unit mass. Shredding mulch residues to small particle size breaks up the continuity of recalcitrant plant tissues, potentially aiding decomposition. This makes the mulch easier to decompose by increasing accessibility and surface area available to soil microbes [23]. Residue size interacts with the C:N ratio and the availability of N in the soil. For example, the decomposition of *short* vs *long residues* may not be accelerated, despite higher contact with soil, when residues are rich in N [24]. The particle size of straw residues (high C:N ratio) did not play an essential role in N immobilization and soil microbial biomass in field conditions with low soil inorganic available N [25]. The immobilization, mineralization, and denitrification of N were higher in shredded residues with high C:N ratio in N rich soil, but only in the short term [23,24]. This suggests that residue particle size does not affect the final amount of nitrogen that is released in the soil but only the timing of the release and availability to plants.

This study aims to improve our understanding of the optimal size of mulch fragments (whole plant versus shredded) and the importance of plant residue diversity. We hypothesise that (i) *long residues* maintain higher fertilization capacity than *short residues* at the end of the growing season, (ii) soil nutrient availability at the later stages of decomposition are affected by both residue size and residue quality (residue C:N ratio, N, and recalcitrant substances), (iii) AMF root colonization increases with increasing residue species richness, (iv) and crop quality is affected by both residue size and residue quality but with no deleterious effect on crop yield.

Our results are discussed in the context of long-term soil health and fertility, with the view of informing the optimisation of long-term mulch use.

## Methodology

### Rhizotron setup

The experiment was conducted at the Crop and Environment Laboratory of the University of Reading in the UK, from June to November 2018. Thirty-six minirhizotrons were constructed from 0.5 cm thick PVC sheeting, each representing an independent replicate (Fig. 1). One side of each rhizotron was clear acrylic to allow observation of root growth and was removable to allow soil sampling. Between observations and sampling, the clear acrylic was covered with thermawrap silver foil to avoid light penetration and to minimize temperature fluctuation. Each rhizotron was 1 m tall to allow root system expansion at depth, and 30 x 5 cm wide, thus providing soil volume of 0.015 m^3^. A layer of gravel c.1.5 cm thick was placed at the bottom to allow drainage, the rest was filled with c.20kg of commercially supplied well-homogenized loamy sand topsoil (bulk soil) with pH 7.3 ± 0,032 SD and sieved to 8 mm. The rhizotrons were kept at 70° angle for the duration of the experiment, with the transparent side facing down so that the roots grow along the transparent side for better observation. Initially, they were placed outside and moved into a greenhouse in October to aid plant senescence.

**Fig. 1.**
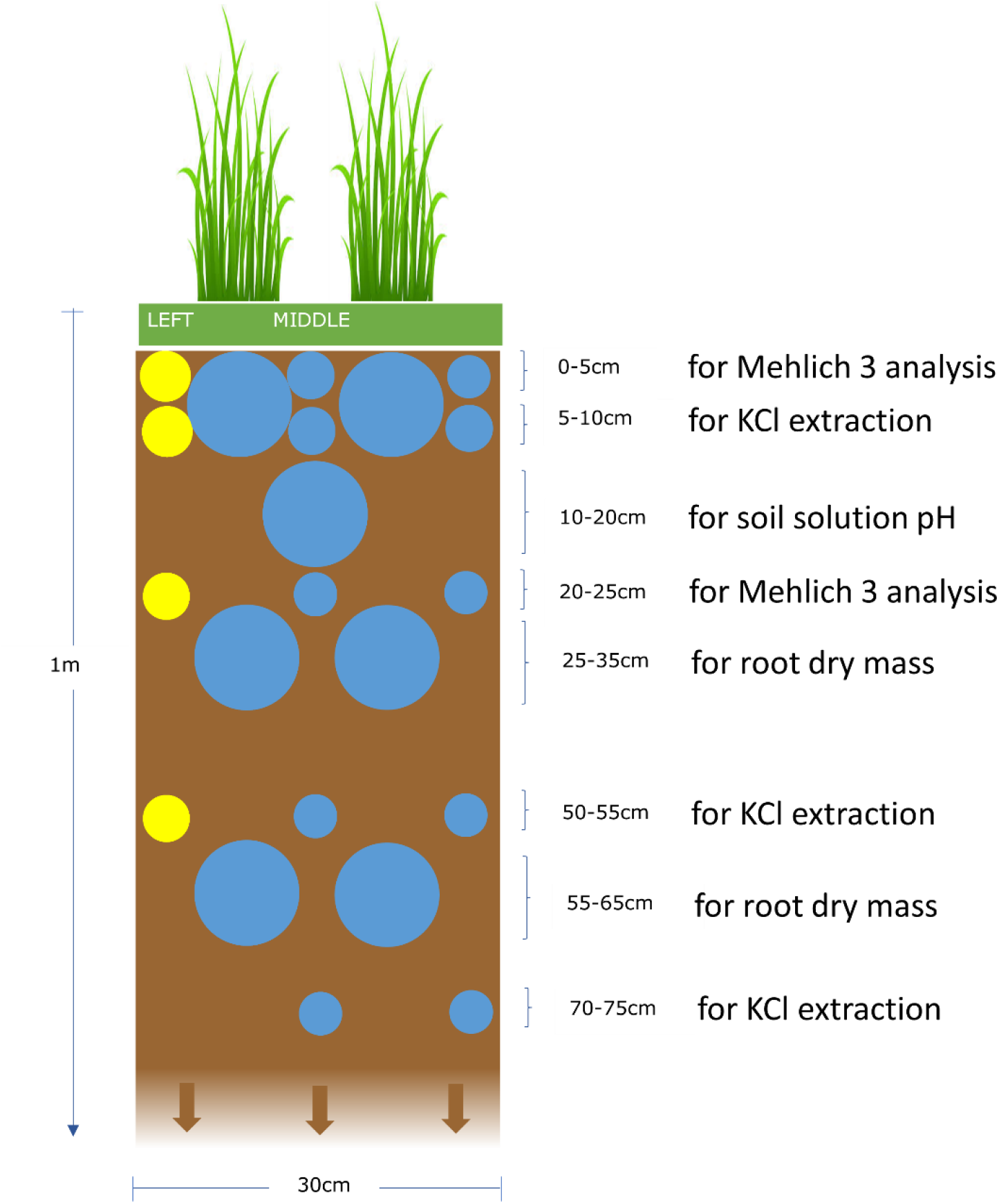
Rhizotron and soil sampling design. Large circles indicate root extraction samples, except for the 10-20 cm depth which was used to measure soil solution pH and bulk density. The large circles at 0-10 cm depth were used to detect AMF colonization in barley roots. Yellow highlighted samples were taken on day 70, blue highlighted on day 137 after mulch application

### Experimental design

Spring barley (*Hordeum vulgare L., var. Laureate*) seeds (56 mg to 66 mg weight) were sown on 11^th^ June 2018 into a germination tray. Plant emergence occurred after 15 days, vigorous plants between 15 cm and 22 cm height were then transplanted into rhizotrons 17 days later. Two plants per rhizotron were planted 15 cm apart and 7.5 cm from each edge (Fig. S1). An automated irrigation system with four drippers per rhizotron was installed to maintain moisture at 55-60% Water Holding Capacity (WHC). Irrigation stopped 106 days after plant emergence to promote seed maturity. Growth stages of the barley plant were observed every 1 to 2 weeks, and the plants were harvested 121 days after mulch application.

Eight biomass residue treatments were established in a fully factorial design (n=4), comprising four plant diversity mixtures (types) and two plant residue particle sizes (*short* of 1.5 cm *(S)*, and *long* of 30cm *(L)* so long as the length of the soil surface in a rhizotron). A control *(C)* with no mulch application was also established. The residue types were: Perennial ryegrass *(**P**)* (1 plant species), Smart Grass™ *(**S**)* (6 species), Biomix™ *(**B**)* (12 species), and Herbal™ *(**H**)* (17 species) (Table S1). No threatened species were used in this research, in accordance with IUCN policy statement on research involving species at risk of extinction. Experimental research including the collection of plant material was conducted on cultivated plants as cover crop plants, in compliance with relevant institutional, national, and international guidelines and legislation. Plant material was source from the University of Reading farm, specifically the DiverseForages grassland field sites. These residue types were suitable to address our hypothesis because they included residue mixtures with different characteristics. Residue characteristics include both residue diversity and functional traits. Residue diversity may concern either the species composition or the species richness which is the number of species participating in a residue mixture. The functional traits of residues may concern either the chemical composition, which determines the residue quality, or the morphological features of residues [26,27]. Fresh plant residue was collected from field plots planted with the aforementioned forage mixtures on 29^th^ June 2018, cut to specified sizes, and placed on the soil surface in rhizotrons randomly allocated to each treatment 4 days later. Plant mass representative of 23g dry mass was used in each treatment. Thus, the 9 treatments were: ***H**L, **H**S, **B**L, **B**S, **S**L, **S**S, **P**L, **P**S, and **C***. No additional fertilization or pesticide treatment was applied to the rhizotrons during the experiment. The encoding *P_(1)_, S_(6)_, B_(12)_*, and *H_(17)_* as well as *PL_(1)_, PS_(1)_ SL_(6)_, SS_(6)_, BL_(12)_, BS_(12)_, HL_(17)_*, and *HS_(17)_* has been adopted where it was considered necessary to indicate the number of species participating in the different residue types or treatments, respectively.

### Soil and plant material sampling and analyses

Soil samples were taken from rhizotrons on two occasions: 70 and 137 days after mulch application (Fig. 1). In total, 21 soil samples were collected from each rhizotron (4 in the first sampling period and 17 in the second), making a total of 756 from all rhizotrons (Fig. S2). All samples, except those with roots, were air-dried and then sieved to <2mm and stored at room temperature. Soil samples with roots were stored at 4°C until root extraction. Roots were extracted by submerging soil samples in tap water over a 1 mm sieve and collecting all floating roots.

After the final harvest, mulch residue, aboveground and belowground barley plant biomass, and soil samples were analyzed to establish key physical and chemical properties of investigated plant-soil systems. Mulch residue and barley biomass were dried for 48 to 72 hours at 80°C until constant weight and then at 105°C for 24 hours to establish dry weight. Total carbon, nitrogen, and protein content of mulch, plant tissue, and seeds, together with total carbon and nitrogen of soil sub-samples, were determined through combustion (LECO CHN 628 analyser, LECO Corporation) [28]. ANKOM 200 Fibre Analyser (ANKOM Technology) [29] was used to measure percentage content of neutral detergent fibre (NDF) (Hemicellulose, Cellulose, and Lignin), acid detergent fibre (ADF) (Cellulose and Lignin), and acid detergent lignin (ADL) (Lignin) of mulch residues according to ANKOM Technology protocols. We then estimated the % Cellulose by subtracting the % ADL from % ADF, and the % Hemicellulose by subtracting the % ADF from the% NDF. Samples were first dried at 80°C to constant weight and ground in a Fritsch grinder (Glen Creston Ltd) with 1mm sieve. Arbuscular mycorrhizal fungi (AMF) colonization and abundance was estimated by black ink staining according to Vierheilig et al. [30].

Air-dried 10 g soil samples sieved to <2 mm were suspended in centrifuge tubes in 25 mL of ultra-pure water and shaken for 15 minutes on an end-over-end shaker to measure soil solution pH [31]. Available N was estimated by KCl extraction method according to Great Britain M.A.F.F. [32] standard protocol using 40 g of air-dry soil in 200 ml 1M potassium chloride solution, measured colorimetrically by a San Continuous Flow Injection Analyzer (SCALAR Instruments) [33]. The Mehlich 3 method [34] was used to evaluate the availability of P, K, Mg, Mn, Fe, Cu and Zn nutrients by Perkin Elmer – Optima 7300 DV ICP-OES analyser (PerkinElmer, Inc.) [35–37]. Air dry soil samples of 2 g and 20 ml of Mehlich 3 extracting solution were used. Other soil samples were air-dried and sieved to <2mm to establish soil texture, which was measured with a hydrometer by adding 50 ml sodium hexametaphosphate solution (SHMP) at 50 g/l to 40 g soil subsamples [38]. Soil water holding capacity (WHC) was estimated gravimetrically using 50 g fresh soil samples with the method described by Harding and Ross [39]. Three samples were taken from the middle of three rhizotrons in random from the depth of 12-16 cm to measure average bulk density [40]. The height of the main stem of barley plants from the soil surface to the base of the flag leaf was used as a proxy of growth rate (cm/d).

#### Statistical analysis

All statistical analyses were carried out in Minitab 19 (Minitab, LLC) [41] except Principal Component Analysis which was conducted in R-Studio (RStudio, PBC) [42]. All measurements from a rhizotron were averaged to obtain a single mean value, the unit of replication of this study being the rhizotron (n=4), and α < 0.05 was used to denote significance. One-way ANOVA and a General Linear Model were used to detect differences between treatments in the case of one factor (treatments, residue type, or residue size) and two factors (type and size of residues), respectively. Measurements repeated in time (day 70, and day 137) or position (middle and right side of rhizotrons) or soil depth were analyzed by Mixed Effects Model with rhizotrons as random factor and type of residues, size of residues, time, depth, and position as fixed factors. All data subjected to analyses of variance were tested for normality and homogeneity, using Darling-Anderson and Levene tests, respectively. When these conditions were not satisfied, data were transformed by log10 or Box-Cox transformation with optimal or rounded λ. Kruskal-Wallis test was conducted instead of one-way ANOVA if transformations did not normalize data variance sufficiently. When a significant treatment effect was observed, a Tukey post-hoc test was also conducted. Comparisons with the control treatment were made using the Dunnett test. Data were separated and analyzed with ANOVA for a specific sampling time or depth, when necessary, where normality or equality of variances were not satisfied even after data transformation. Descriptive statistics included means and standard deviations rather than standard errors unless it was otherwise stated.

Simple regression analysis was conducted in order to examine the significance and the degree of the effect of a single independent variable to a response variable in cases where there was an apparent influence (residue dry mass loss vs initial residue NDF or N content or initial residue C:N ratio, soil K vs initial residue C:N ratio, AMF root colonization vs residue species richness, seed protein vs main stem elongation rate as continuous predictor variable and residue size as categorical predictor variable). Homogeneity of variance of the data was tested with Levene test, and normality of data distribution with Anderson-Darling test. The Spearman correlation was used when normality or equality of variances of data were not satisfied. Pearson correlation was used to assess the significant correlations between variables.

Multivariate analysis was also conducted on data collected from barley plants and soils using Principal Component Analyses (PCAs) to assess differences between treatments in several variables simultaneously, concerning measurements on barley plants or soil nutrients, and the relations of the treatments with those variables.

## Results

### Residue dry mass loss, and initial and final quality

Residue (mulch) dry mass loss due to decomposition was significantly different between treatments (N = 4, F = 2.57, p-value = 0.040) and specifically between *BS_(12)_* (71.71 ±7.21) and *PL_(1)_* (47.78 ±6.44) (T-value = −3.40, p-value = 0.041, Fig. 2).

**Fig. 2.**
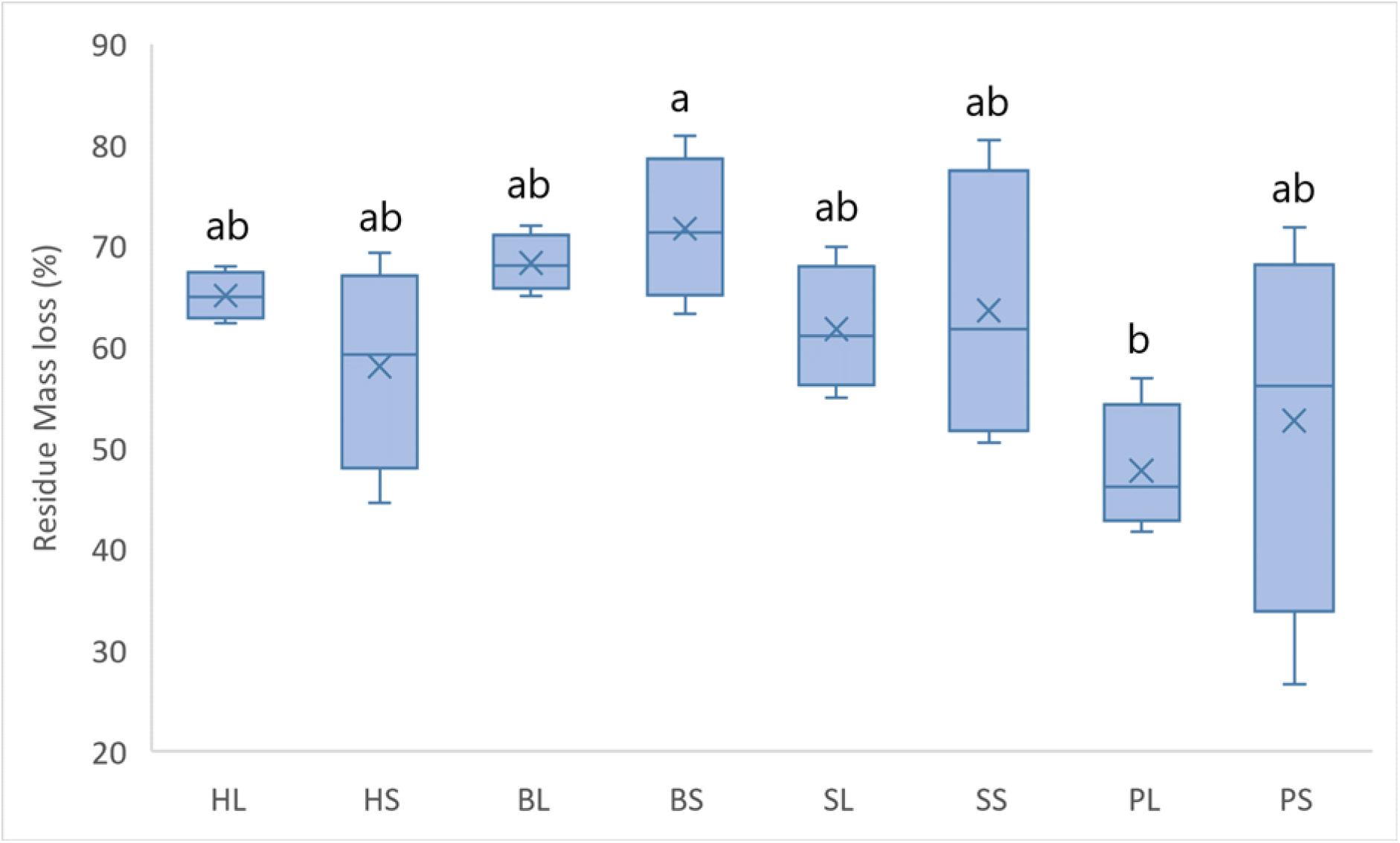
Final residue dry mass loss (121 days after mulch application). Mean values are depicted with x (N = 4, F = 2.57, p-value = 0.040). The residue types were: Perennial ryegrass *(**P**)* (1 plant species), Smart Grass *(**S**)* (6 species), Biomix *(**B**)* (12 species), and Herbal *(**H**)* (17 species). Residue treatments including four plant diversity mixtures (types) and two plant residue fibre sizes (*short* of 1.5 cm *(S)*, and *long* of 30cm *(L)*). Treatments that do not share a common letter are significantly different (p < 0.05)

Further analysis showed a significant effect of residue diversity (residue types) on residue dry mass loss (N = 8, F = 5.40, p-value = 0.006). There were significant differences between *P_(1)_* (50.26 ±13.33) and *B_(12)_* (70.02 ±5.39) (T-value = −3.98, adjusted p-value = 0.003), but no significant between *long* and *short* residue size (N = 16, F = 0.05, p-value = 0.823).

The *P_(1)_* residue type had the lowest initial C:N ratio (24.2 ±2.268) while *S_(6)_* had the highest (33.5 ±1.65), *B_(12)_* had 30.4 ±3.64 and *H_(17)_* 28.1 ±1.801 (N = 8, F = 20.24, p-value < 0.001, Tables S2 and S3). Type *P_(1)_* had the highest mean value (41.644 ±0.160) of initial C content of residues, significantly different from *B_(12)_* (41.229 ±0.256), and *H_(17)_* (41.207 ±0.195) (N = 8, p-value = 0.001, Tables S2, and S3). *S_(6)_* type had the lowest initial N content (1.238 ±0.055) while *P_(1)_* had the highest (1.734 ±0.170), and there were significant differences between types (N = 8, F = 19.55, p-value < 0.001, tables S2 and S3). The same pattern was observed in initial protein content of residues (N = 8, F = 19.55, p-value < 0.001). There were no significant differences in initial ADL (lignin) content between residue types (N = 4, Kruskal-Wallis test p-value = 0.277, Fig. S3, Table S4). In addition both NDF (hemicellulose + cellulose + lignin) and ADF (cellulose + lignin) contents include lignin. There was a significant positive correlation between NDF and ADF (r = 0.938, p-value < 0.001) and no significant correlation between NDF and lignin (r = −0.021, p-value = 0.937) or between ADF and lignin (r = −0.058, p-value = 0.832). Moreover, differences between residue types in ADF were statistically more significant (N = 4, F = 5.76, p-value = 0.011) than in cellulose alone (Kruskal Wallis test, N = 4, H-value = 6.60, p-value = 0.086), while it was of equal significance in both NDF (N = 4, F = 16.14, p-value < 0.001) and in hemicellulose alone (N = 4, F = 23.04, p-value = < 0.001) (Fig. S3, Tables S4 and S5). For these reasons, NDF was used in the experiment as a measure of residue recalcitrance.

Residues harvested at the end of the experiment showed significant differences in C:N ratio between treatments (N = 4, F = 4.27, p-value = 0.003). *PL_(1)_* treatment which had the highest mean value was significantly higher than *HS_(17)_* and *BS_(12)_* (Fig. 3a, Tables S6 and S7). Further analysis showed significant differences in final C:N ratio between types (N = 8, F = 7.57, p-value = 0.001) as well as between *short* (15.029 ±2.099) and *long* (16.744 ±2.595) residues (N = 16, F = 6.66, p-value = 0.016). The C:N ratio was significantly higher in *P_(1)_* (17.959 ±2.407) than in *B_(12)_* type (13.883 ±1.797) (T-value = 4.33, adjusted p-value = 0.001), in *P_(1)_* than in *H_(17)_* type (14.922 ±2.000) (T-value = 3.23, adjusted p-value = 0.018), and in *S_(6)_* (16.786 ±1.655) than in *B_(12)_* type (T-value = 3.09, adjusted p-value = 0.024). Final C content of residues was also significantly higher (59.20%) in *long* residues (30.95 ±4.68) than *short* residues (19.44 ±5.89) (N = 16, F = 36.74, p-value < 0.001). *PL_(1)_, SL_(6)_*, and *HL_(17)_* were significantly higher than *BS_(12)_, HS_(17)_*, and *PS_(1)_* (N = 4, F = 6.02, p-value < 0.001, Fig. 3b, Tables S6 and S7). There were significantly higher final N contents in *HL_(17)_, BL_(12)_, SL_(6)_* treatments than in *HS_(17)_*, and *PS_(12)_* (N = 4, F = 5.68, p-value = 0.001, Fig. 3c, Tables S6 and S7). In addition, final N content of residues was significantly higher (44.06%) in *long residues* (1.854 ±0.154) than *short* ones (1.287 ±0.303) (N = 16, F = 38.05, p-value < 0.001).

**Fig. 3.**
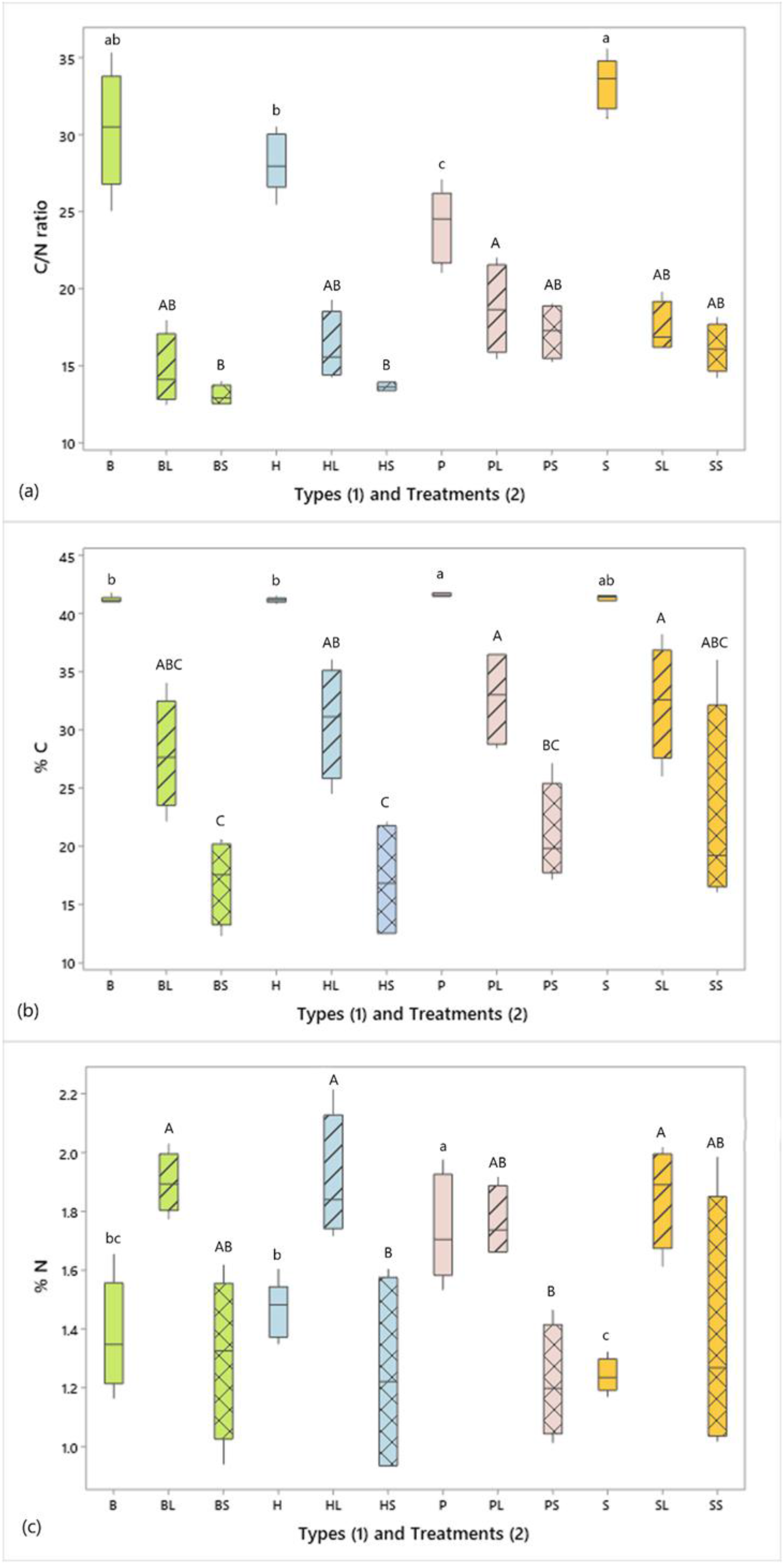
Box and whiskers plots of plant residue initial and final (a) C:N, (b) % C, and (c) % N ratio for the different types (*B, H, P, and S*) (initial residues) and Treatments (*BL, BS, HL, HS, PL, PS, SL, and SS*) (final residues). The residue types were: Perennial ryegrass *(**P**)* (1 plant species), Smart Grass *(**S**)* (6 species), Biomix *(**B**)* (12 species), and Herbal *(**H**)* (17 species). Residue treatments including four plant diversity mixtures (types) and two plant residue fibre sizes (*short* of 1.5 cm *(S)*, and *long* of 30cm *(L)*). Lower case letters refer to comparisons between initial residue types. Upper case letters refer to comparisons between final residue treatments. Mean values that do not share a common letter are significantly different (p < 0.05)

There were no statistically significant correlations between C and N content in final *long* residues: *HL_(17)_* r = 0.518 (p = 0.482), *BL_(12)_* r = 0.461 (p = 0.539), *SL_(6)_* r = 0.801 (p = 0.199), *PL_(1)_* r = −0.199 (p = 0.801). In contrast, there were significant positive correlations between C and N content in final *short* residues in all treatments except in *PS_(1)_* treatment: *HS_(17)_* r = 0.999 (p = 0.001), *BS_(12)_* r = 0.970 (p = 0.030), *SS_(6)_* r = 0.970 (p = 0.030), and *PS_(1)_* r = 0.869 (p = 0.131). The same was true considering all *long* residue (r = 0.260, p = 0.331) and all *short* residue (r = 0.875, p < 0.001) treatments together.

Simple linear regression analysis was conducted using the residue dry mass loss as response variable (y) and either the initial NDF content or the initial N content or the initial C:N ratio of residues (x) as predictor variable (N = 8). The analysis showed that the regression model could explain only r^2^ = 21.12% or 24.50% or 21.36%, respectively of the variation of the response variable. However, the influence of the predictor was statistically significant in all cases (p-value = 0.008 for NDF, 0.004 for N, and 0.008 for C:N ratio). The regression equations were y = 113.1 - 1.190x for NDF, y = 106.2 - 30.99x for N, and y = 16.2 + 1.548x for C:N ratio.

### Soil nutrient content

Soil samples taken at harvest time on day 137 from 10-20 cm depth from the middle of the rhizotrons showed no significant differences in soil solution pH between treatments (N = 4, F = 0.40, p-value = 0.909).

The effect of residue diversity and of residue size on soil NO_3_^-^ content was not significant on day 70 at 50-55cm depth (the only timepoint and depth that they were detected), (N = 4, F = 0.46 and p-value = 0.712 for type, F = 0.30 and p-value = 0.591 for size). However, mean value of all treatments (1.407 ±0.852) was significantly lower than that of bulk soil (initial soil, prior to its use in rhizotrons) (11.334 ±1.533, N = 4, F = 40.80, p-value < 0.001).

Soil NH_4_^+^ content was highly significantly lower on day 70 (1.302 ±0.356) than day 137 (1.678 ±0.379) (N = 64, F = 46.65, p-value < 0.001) and significantly higher at the top 5-10 cm (1.689 ±0.280) than at 50-55 cm (1.292 ±0.428) depth (N = 64, F = 52.13, p-value < 0.001, Tables S8, S9, and Fig. S4).

Depth significantly affected all soil nutrient contents (P, K, Mg, Fe, Mn, Zn, and Cu), and time significantly affected all nutrients except Fe and Mn (Tables S10 and S11). Macronutrients P and Mg had higher mean values at 20-25 cm depth than at 0-5 cm, but the opposite was true for K. Macronutrient concentrations were higher on day 70 than on day 137 at both 0-5 and 20-25 cm depths. Further analysis showed significant differences in soil K content between diverse residue types on day 137 at 0-5 cm depth (N = 8, F = 4.16, p-value = 0.017, Fig. 4). Tukey’s post-hoc testing showed significant differences between *S_(6)_* (mean = 102.60 ±31.70) and *P_(1)_* (68.66 ±13.11) (T-value = 2.92, p-value = 0.035), and between *S_(6)_* and *B_(12)_* (68.44 ±24.12) (T-value = 3.17, p-value = 0.020) types. Spearman’s correlation confirmed there was a positive and statistically significant association between the soil K content and the initial C:N ratio of residues (Spearman ρ = 0.439, ρ = 0.007).

**Fig. 4.**
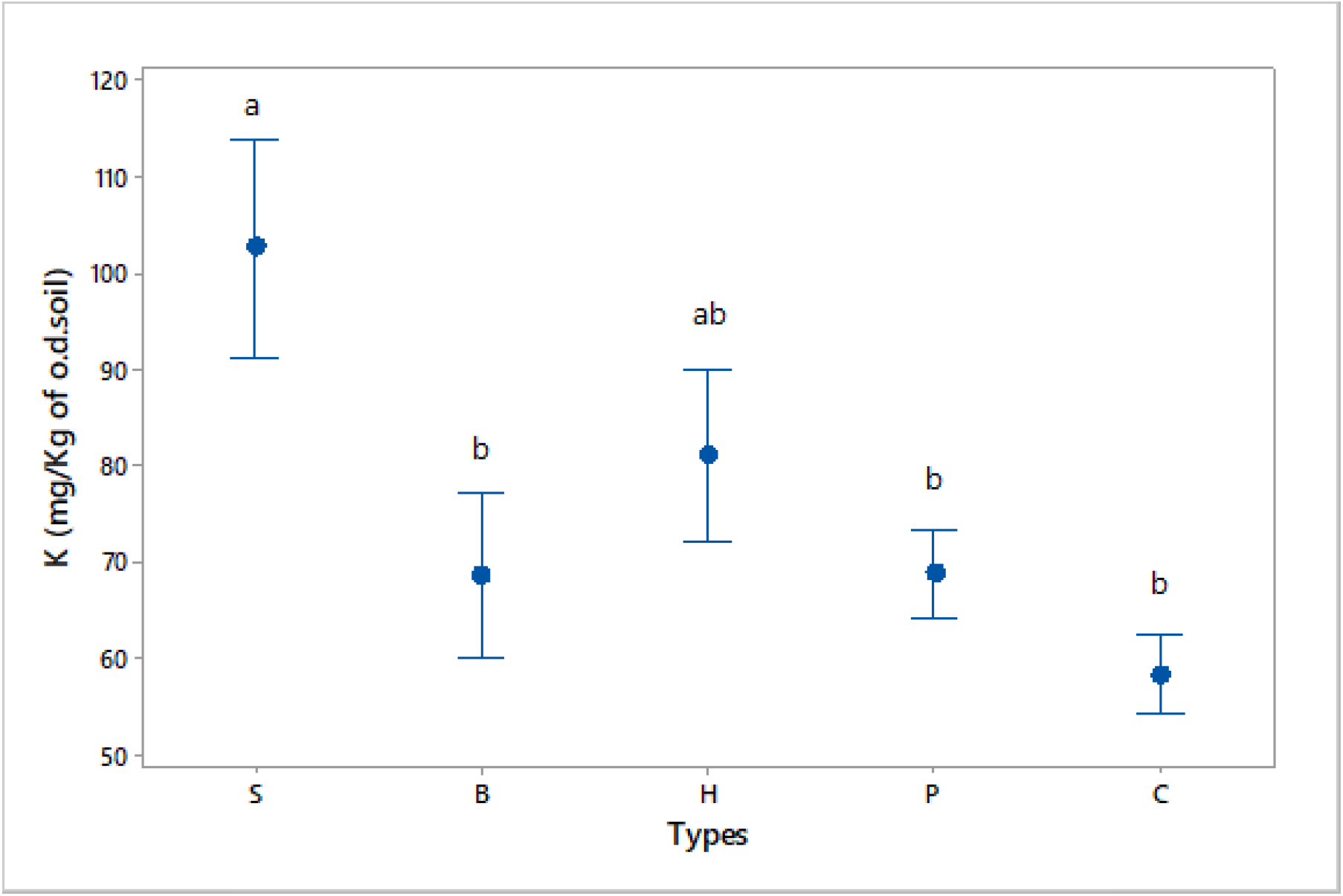
The effect of different types (*S, B, H, P* in descending order of their initial C:N ratio) of residues on soil K concentration (Mehlich 3 extraction), in comparison to unamended Control *(C)*, 137 days after mulch application. Results include all types, and depth of 0-5 cm (N = 8). The residue types were: Perennial ryegrass *(**P**)* (1 plant species), Smart Grass *(**S**)* (6 species), Biomix *(**B**)* (12 species), and Herbal *(**H**)* (17 species). Means and bars of one standard error from the mean are depicted. Types that do not share a common letter are significantly different (p-value < 0.05)

All micronutrients (Fe, Mn, Zn, and Cu) had significantly higher mean values (p-values < 0.001) at 0-5 cm depth than at 20-25 cm. Further analysis revealed significantly lower soil Zn content (−38.30%) in *long* residues (21.73 ±8.77) than *short* (35.22 ±15.69) on day 137 at 0-5 cm depth (N = 16, F = 7.53, p-value = 0.011).

The influence of the different treatments on soil nutrients (P, K, Mg, Fe, Mn, Zn, and Cu) on a multivariate basis, at the depths of 0-5 cm and 20-25 cm, was tested in Principal Component Analyses on day 70 (Fig. S5), and on day 137 after mulch application (Fig. S6). Although in most cases treatments were overlapping, *Control* seemed to demonstrate the most negative relation with soil nutrients in comparison to the other treatments, most clearly at 0-5 cm depth on day 70 (Fig. S5a). Also the contribution of *S_(6)_* type in enrichment of soil K in comparison to the other treatments was obvious at 0-5 cm depth on day 137 (Fig. S6a).

Cultivation of barley resulted in statistically significant reduction in soil K (N = 4, Kruskal-Wallis p-value < 0.001) and Mn (N = 4, F = 2.50, p-value = 0.011) in comparison to the bulk soil.

Samples taken from the middle of the rhizotrons, the area of plant root interaction, on day 137 revealed significant differences in soil nutrient contents in comparison to samples from the right side of rhizotrons, considering both samples from 0-5 and 20-25 cm depths (N = 64, F = 16.27, p-value < 0.001 for soil K, F = 56.91, p-value < 0.001 for P, F = 6.60, p-value = 0.012 for soil Mn, F = 14.52, p-value < 0.001 for soil Zn, and F = 16.51, p-value < 0.001 for soil Cu). Therefore, the effect of plant root interaction should be considered in experiments.

### Barley plants

Residue size significantly affected the length of ears (N = 32, F = 4.96 and p-value = 0.030). Treatments with *short* size residues had higher mean values (8.337 ±1.453) than those with *long* size (7.778 ± 0.052). Total yield was not significantly affected by the different treatments (N = 8, F = 0.29, p-value = 0.968). Likewise, barley seed protein content was not statistically significantly different between treatments of long and short residues (N = 4, F = 1.90, p-value = 0.182) or between treatments of different types (N = 4, F = 1.50, p-value = 0.242). However, in all types except *P_(1)_* all treatments with long size residues had higher values of protein content than treatments of the same type with short size residues, and Control had lower value than any treatment with long residues (Table S12). Spearman correlation showed a significant negative correlation (Spearman ρ = −0.529, p-value = 0.001) between length of ear and seed protein content (%).

The Arbuscular Mycorrhizal Fungi (AMF) root colonization, detected on day 137, showed no significant differences as a result of treatments (*HL, HS, BL, BS, SL, SS, PL, PS, C*) (N = 4, F = 1.16, p-value = 0.356) or residue size (N = 16, F = 0.09, p-value = 0.772). However, mean values indicated that residues of higher species richness had higher AMF colonization. *Control* had the lowest mean value (Table 2).

Linear regression analysis showed that residue species richness could explain only r^2^ = 15% of the AMF root colonization variation. However, the influence of the predictor was statistically significant (N = 8, p-value = 0.02). The regression equation was y = 30.60 + 0.365x, which means that for every plant species that is added in the residue mixture an increase of 0.365% in AMF root colonization was expected (Fig. S7).

### Barley plant growth rate

All treatments followed the same pattern of stem elongation (Fig. 5).

**Fig. 5.**
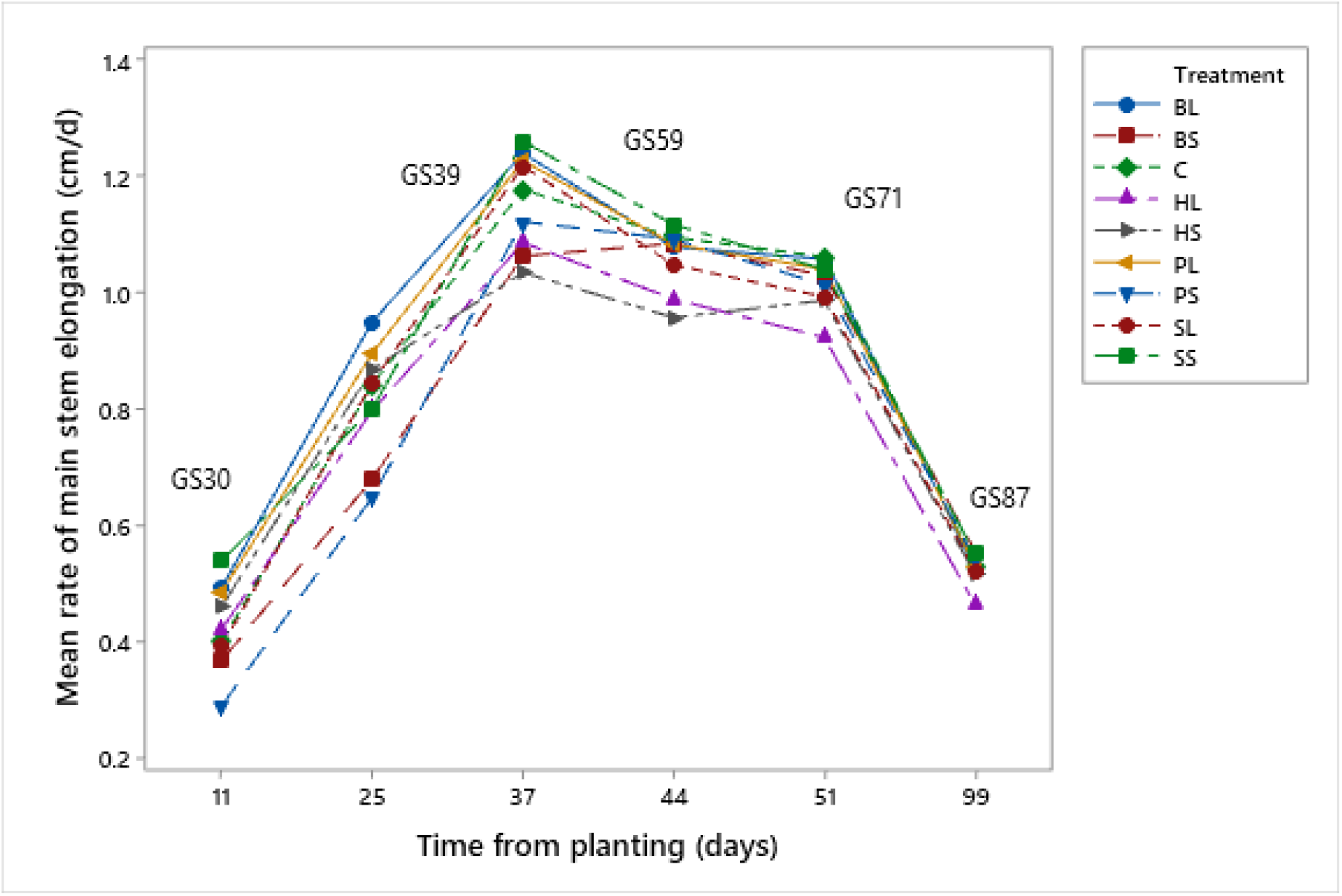
Mean rate of main stem elongation (cm/d) of barley plants from soil surface to the base of the flag leaf for the different treatments (*BL, BS, C, HL, HS, PL, PS, SL, SS*, and *C*), 11, 25, 37, 44, 51, and 99 days after planting (28, 42, 54, 61, 68, and 116 days after plant emergence). GS = growing stage [43]. The residue types were: Perennial ryegrass *(**P**)* (1 plant species), Smart Grass *(**S**)* (6 species), Biomix *(**B**)* (12 species), and Herbal *(**H**)* (17 species). Residue treatments including four plant diversity mixtures (types) and two plant residue fibre sizes (*short* of 1.5 cm *(S)*, and *long* of 30cm *(L)*). Control treatment *(C)* was with no residues

Main stem elongation rate (cm/d) was significantly higher in *long* residues (0.871 ±0.230) than *short* residues (0.749 ±0.257) 25 days after planting (42 days after plant emergence) (N = 16, F = 4.00, p-value = 0.050). However, 99 days after planting (116 days after plant emergence) main stem elongation in *long* residue treatments were significantly lower (0.509 ±0.061) than in *short* ones (0.540 ±0.0521) (N = 16, F = 4.98, p-value = 0.003). Contrary, residue type did not affect significantly main stem elongation rate.

Linear regression analysis of seed protein content (%) as response variable, main stem elongation rate (MSER) as continuous predictor variable, and residue size (*long, short*, and *Control*) as categorical predictor variable on day 25 after planting was conducted (Fig. S8). The analysis showed that main stem elongation rate could explain only r^2^ = 15.13% of the final seed protein content, but the influence was significant (N = 8, F = 4.62, p-value = 0.040). The regression equations were:

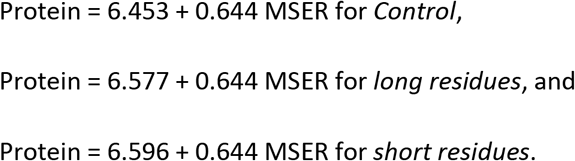

The influence of the different treatments on several variables concerning barley plant biomass (variables v1 and v2), the relation of barley plants with symbiotic microbes (variable v3), and seed quality and yield (variables v4 to v6) on a multivariate basis, was tested in a Principal Component Analysis (Fig. 6).

**Fig. 6.**
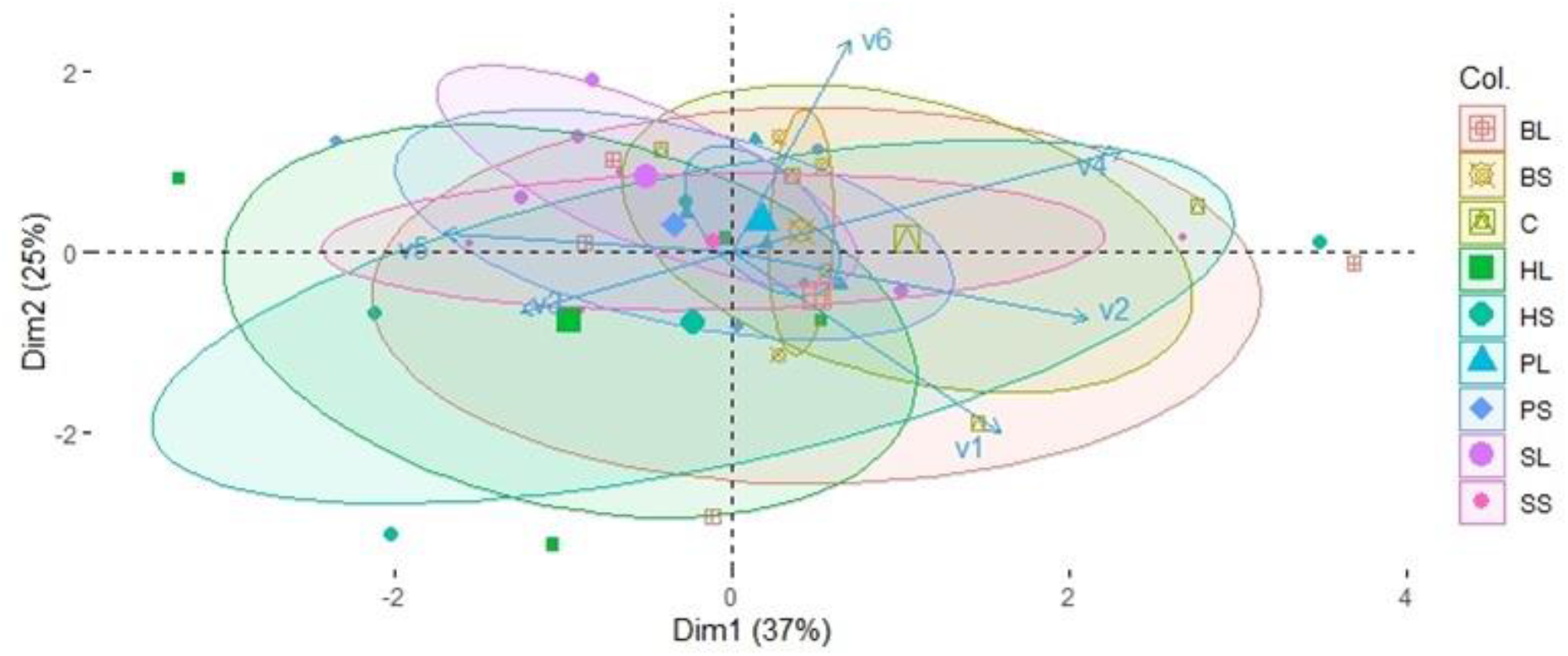
PCA ordination of variables v1 = Barley root dry mass (g) (from combined samples at 25-35cm and 55-65 cm depth), v2 = barley plant dry mass (g) per plant per rhizotron (without ears and roots), v3 = % AMF colonization in barley roots, v4 = total seed dry mass (g) per plant per rhizotron, v5 = % barley seed protein content, v6 = % barley seed carbon content, for the treatments *HL, HS, BL, BS, SL, SS, PL, PS*, and *Control (C)* concerning measurements on barley plants. The residue types were: Perennial ryegrass *(**P**)* (1 plant species), Smart Grass *(**S**)* (6 species), Biomix *(**B**)* (12 species), and Herbal *(**H**)* (17 species). Residue treatments including four plant diversity mixtures (types) and two plant residue fibre sizes (*short* of 1.5 cm *(S)*, and *long* of 30cm *(L)*). Control treatment *(C)* was with no residues. Large points depict mean values, while ellipses depict confidence intervals (α = 0.05) of mean values for each treatment. The percentages represent data variation explained by the two first Principal Components (Dim1 and Dim2), bottom axis represents Dim1 normalized score, and left axis represents Dim2 normalized score

In Fig. 6 it is evident, *HL_(17)_* treatment had the strongest positive relationship with AMF colonization (v3 variable) and the strongest negative with total seed dry mass (v4), opposite *Control* treatment. In the rest of variables differences between treatments were not clear confirming the lack of significant differences as treatments were highly overlapped.

## Discussion

### Residue quality and degree of decomposition

The initial C:N ratios of the residue types were all significantly different, ranging between 15 and 33. Soil decomposer microbes have a C:N ratio around 8 in their body and assimilate about 1/3 of the decomposed C. Therefore, an initial residue C:N ratio ≤24 is considered to drive to net N mineralization even at later stages of decomposition contrary to residues with C:N ratio >24 [44]. However, this threshold may be risen to more than 30 by the activity of soil fauna, but normally <35 [45,46]. C:N ratio affects organic matter decomposition in N-poor soils, while its effect in N-rich soils can be insignificant [47]. Apart from C:N ratio, decomposition is greatly affected by residue recalcitrance as well [48]. Interestingly, *P_(1)_* type had the lowest C:N ratio but the highest recalcitrance, not significantly different from that of *S_(6)_* type despite *S_(6)_* having the highest C:N ratio (Table 1).

**Table 1.** Classification of residue types (*H, B, S, and P*) and treatments (*HL, HS, BL, BS, SL, SS, PL, PS*, and *C*) in descending order of their initial or final mean values according to different properties. The residue types were: Perennial ryegrass *(**P**)* (1 plant species), Smart Grass *(**S**)* (6 species), Biomix *(**B**)* (12 species), and Herbal *(**H**)* (17 species). Residue treatments including four plant diversity mixtures (types) and two plant residue fibre sizes (*short* of 1.5 cm *(S)*, and *long* of 30cm *(L)*). Control treatment *(C)* was with no residues. NDF = Neutral Detergent Fiber (Hemicellulose, Cellulose and Lignin), ADF = Acid Detergent Fiber (Cellulose and Lignin), and ADL = Acid Detergent Lignin (Lignin)

| Properties | Residue types | Properties | Treatments |
| --- | --- | --- | --- |
| Initial C:N ratio | $S^a > B^{ab} > H^b > P^c$ | Final C:N ratio | $PL^a > SL^{ab} > PS^{ab} > HL^{ab} > SS^{ab} > BL^{ab} > HS^b > BS^b$ |
| Initial % C | $P^a > S^{ab} > B^b > H^b$ | Final % C | $PL^a > SL^a > HL^{ab} > BL^{abc} > SS^{abc} > PS^{bc} > HS^c > BS^c$ |
| Initial % N | $P^a > H^b > B^{bc} > S^c$ | Final % N | $HL^a > BL^a > SL^a > PL^{ab} > SS^{ab} > BS^{ab} > HS^b > PS^b$ |
| % initial NDF | $P^a > S^b > B^{bc} > H^c$ | | |
| % initial ADF | $P^a > S^{ab} > B^b > H^b$ | | |
| % initial ADL | $S > H > P > B$ | | |
| Diversity | $H > B > S > P$ | | |

*Types or treatments of the same property that do not share a common letter are significantly different ( $p < 0.05$ ).*

**Table 2.** Mean values and standard error of AMF root colonization (%) in all treatments (*HL, HS, BL, BS, SL, SS, PL, PS*, and *C*) at harvest time (on day 137 after mulch application). The residue types were: Perennial ryegrass *(**P**)* (1 plant species), Smart Grass *(**S**)* (6 species), Biomix *(**B**)* (12 species), and Herbal *(**H**)* (17 species). Residue treatments including four plant diversity mixtures (types) and two plant residue fibre sizes (*short* of 1.5 cm *(S)*, and *long* of 30cm *(L)*). Control treatment *(C)* was with no residues. Mean values and standard errors are included (N = 4)

| **Treatments** | **Mean ±SE** | **Treatments** | **Mean ±SE** | **Treatments** | **Mean ±SE** |
| --- | --- | --- | --- | --- | --- |
| **HL** | **35.52 ±1.67** | **SL** | **37.94 ±3.25** | **C** | **28.84 ±2.34** |
| *HS* | 36.27 ±4.30 | *SS* | 32.42 ±2.70 |  |  |
| **BL** | **34.88 ±3.44** | **PL** | **29.35 ±1.54** |  |  |
| *BS* | 35.41 ±2.33 | *PS* | 31.05 ±3.99 |  |  |

In all cases, treatments with *long residues* resulted in higher final C:N ratio, C, and N content than those of the same type with *short residues* at the end of the growing season. Higher final C:N ratio in residues of the same type denotes lower decomposition rate, because C:N ratio is decreasing during decomposition after an initial short increase [49]. Considering the dry mass loss of residues was not significantly affected by their size, this is clear evidence that *long residues* contributed to a higher accumulation of C and N on soil surface, in the form of C and N content retained in residues, at the end of the growing season than the *short residues*. This suggests possible positive effects on the mineralization of C and N, and possibly of other nutrients, for subsequent crops with continuous application of long versus short residue mulches (higher fertilization capacity).

Moreover, the final C:N ratio was highly affected by residue quality. Decomposition drive to lowering of the C:N ratios of the different residue types with a tendency to equalize [50]. However, the lignin and other recalcitrant fractions of residues that were less affected during decomposition [51] resulted in higher final C:N ratios of residues. Indeed, final C:N ratio was higher at *P_(1)_* and *S_(6)_* types and especially at *PL_(1)_* and *SL_(6)_* treatments (*long* residues of higher recalcitrance) than at *B_(12)_* and *H_(17)_* and especially at *BS_(12)_* and *HS_(17)_* (short residues of lower recalcitrance) (Table 1). Recalcitrance was distinguished by initial NDF content since there were no significant differences in initial ADL (lignin) content of the residues. Besides, Jensen et al. [52] found that hemicellulose and cellulose were the factors that explained the greatest variability of C mineralization rather than lignin. Therefore, C and N mineralization of residues were affected mainly by residue size and initial NDF content rather than by initial C:N ratio and lignin content. It is possible that lower final N content in *short residues* was due to slightly increased N availability from more recalcitrant fractions, caused by chopping, and subsequent N uptake by plants or N losses by leaching. Chopped material is decomposed faster during the early days or weeks of decomposition because of larger exposure of the residue surface to direct contact with soil and soil microbes [51]. Therefore, the final residue quality was affected by both residue size and residue quality.

Moreover, the lack of correlation or the weak correlations between final C and N of treatments with *long residues* versus the strong relevant correlations in *short residues* may be an indication that different groups of microbes were responsible for the decomposition in each case. Assuming that the *short residues* are in better contact with the soil, the decomposition of *short residues* should be dominated by bacteria, and those of *long* residues by hyphae-producing fungi [3,53]. Bacteria preferentially decompose the readily available fractions leaving the more recalcitrant lignin fraction, while fungi decompose the more recalcitrant fractions [54]. Thus, the correlation index (r) between C and N of final residues could possibly be used to estimate the degree and the microbial pathway of decomposition of mulch residues.

Dry mass loss was not significantly affected by residue size in the long term, which corresponds to previous findings [24,55]. However, it seems to be related to both residue recalcitrance and C:N ratio or N content. This was confirmed by the simple linear regression analysis where the influences of NDF, N, and C:N ratio on residue mass loss were statistically significant, although each one of the three independent variables could explain only a small part of the variation of the dry mass loss (from 21.12 to 24.50%). Decomposition of lignin in residues typical for high recalcitrance and high N content is restricted because the relevant decomposer microbes need a readily available C source. Excessive N makes these microbes more vulnerable to competition with other microbes [56]. Therefore, the higher the initial NDF content of residues the lower the residue dry mass loss (primarily), and the higher the initial N content in residues of higher recalcitrance the lower the final dry mass loss. Likewise, Xu et al. [57], in a one-year experiment, noticed significant differences in biomass loss, higher in species with higher C:N ratio but low recalcitrance (polyphenol, lignin, and tannin) in residues cut to less than 2 cm and buried at 10 cm depth in sealed litter bags. The microorganisms adapted to the decomposition of recalcitrant substances are favoured by residues with a higher C:N ratio as they are able to outcompete other types of microbes [24]. Alternatively, higher diversity of residues may trigger higher diversity of decomposer microbes and consequently higher specificity in enzymatic activity during decomposition [20], driving to higher residue mass loss. Shu et al. [58] also observed higher microbial activity in residue mulch of higher diversity (mixed than single plant species), but not higher microbial biomass or microbial community composition, and attributed this effect to the activation of dormant microbial populations by residue mixtures.

The fact that long residues resulted in higher final C and N content than short residues, while at the same time residue mass loss was not significantly affected by residue size, implies that long residues maintained higher fertilization capacity than short residues at the end of the growing season, in line with our hypothesis.

### Dynamics of soil nutrients

At and after barley growing stage GS61 [43], the N uptake requirement of barley plants declines as plant growth has almost been completed and plants redistribute N to feed the developing grains [59]. The decomposition of the residues initially affected the availability of NH_4_^+^ and soluble organic N (SON). NH_4_^+^ is typically not prone to leaching [60], but may be subjected to nitrification [61] and the resulting NO2^-^ and NO_3_^-^ which may leach downwards – a process detected in our samples from the 50-55 cm depth on day 70. Subsequently, the nitrification activity was restricted considerably on day 137 probably due to cessation of irrigation to promote seed maturity [62]. Both *short* and *long residues* led to significantly higher values of soil NH_4_^+^ on day 137 than on day 70 (Tables S8, S9, and Fig S4). It is possible, the observed lower values of soil NH_4_^+^ on day 70 to be attributed to plant uptake because maximum nutrient uptake is up to GS70 (end of flowering stage) [63]. Moreover, mean values of *short residues* were higher than those of *long residues* on day 70 and the opposite was true on day 137, although differences were not significant (Table S8). This shows that the rate of N mineralization was initially higher in *short residues*, but it was reversed in favour of long residues sometime up to day 137. This is supported by work from, Angers and Recous [23] who observed initially higher decomposition rates at small particle size residues incorporated into soil, followed by higher rates at long size (up to 10 cm) one. In contrast, experiments with fine and ground (usually <1 cm) particle size residues mixed with soil showed effect of particle size on N dynamics only in early stage of decomposition [24,55,64].

Mean values of soil NO_3_^-^ were not significantly affected by the treatments, although they were far lower than the initial measurements in bulk soil. In addition, no significant effect of residue diversity or functional traits on soil NH_4_^+^ content was observed on both day 70 and day 137. Contrary to our experiment, N mineralization was reported to be affected by residue chemical composition after 100 days of incubation, where high positive correlations had been found between N mineralization and N to lignin content ratio of residues [65]. Any such effect in our experiment possibly either occurred at earlier stages of decomposition and differences were soon compensated or it was negligible. Moreover, Fox et al. [66] concluded that when incorporating residues into soil, (lignin + polyphenol):N ratio could be used as a reliable predictor of N mineralization rate of residues, but they did not find significant correlation between N mineralization and residue N or lignin or polyphenol content alone.

Soil K was higher in all treatments with residues than in *Control*, in concordance with previous research [67]. The return of plant residues to the soil is a considerable source of soil K replenishment. K is readily released as K^+^ ions to soil solution during decomposition because, it remains in plants in ionic form in the cell solution and contributes, like Mg, to the production of extracellular enzymes [68]. We hypothesised that the residue chemical composition (quality) as well as residue size would affect nutrient dynamics. Our data showed that K was the only nutrient that was significantly affected by residue chemistry. Soil K was higher in residues with higher C:N ratio, and this positive correlation was statistically significant as it was confirmed by Spearman’s correlation. Probably, the fact that K largely remains in the plant cells in ionic form makes it less prone to be bound in residue recalcitrant substances and therefore its release to soil solution during decomposition is dependent on initial residue C:N rather than NDF content. It is possible, residues of higher quality released higher amounts of K earlier due to higher initial rates of decomposition, while the opposite was true with lower quality residues [69].

Soil micronutrients Fe, Mn, Zn, and Cu had significantly higher mean values at 0-5 cm depth than at 20-25 cm. Normally their concentrations are higher at surface (Ap) soil horizons [70]. Soil Zn was the only nutrient that was significantly affected by residue size, in confirmation to our hypothesis, with higher values in *short residues*, but likewise K, only on day 137 at 0-5 cm depth. It is estimated that about 50% of soils cultivated with cereals worldwide suffer from low Zn content with negative impact in production and grain quality [71]. Increased release of Zn at later stage of decomposition by *short residues* may be deemed as a precursor of an increased Zn release by *long residues* in a successive cash crop. Contrary, higher soil Zn content could be the result of phytosiderophore exudates by barley plants (graminaceous monocotyledonous species) in Fe or Zn deficient soil [72]. In this case *long residues* provide an advantage by slow and stable release of micronutrients avoiding Zn deficiency.

Principal Component Analysis showed that *Control* was generally a negatively related treatment with all soil nutrients in comparison to the other treatments. This indicates the value of residue mulch in enrichment of soil nutrients even at 20-25 cm depth on both day 70 and day 137.

### Impact on barley plants

Yield of barley plants was not affected by residue diversity or size. This is consistent with earlier studies, e. g. Reichert et al. [73] found no differences in crop (cassava) yield between treatments with chopped mulch, although it concerned residues with very small particle sizes. On the contrary, Awopegba et al. [74] noticed significant differences in crop (maize) yield between types of treatments with chopped material applied on soil with a traditional hoe. In our experiment, there were indications that *long residues* resulted in higher seed protein content than *short residues. Long residues* have slower rate of decomposition and are thus able to provide more N at later stage of decomposition (GS37) [43] than the *short residues*, which is crucial to increase protein content of grains [75]. Indeed, *long residues* resulted in significantly higher rates of barley stem elongation 42 days after plant emergence (at GS31 to GS39 – rapid stem elongation stage) in comparison to *short residues*. This is an indication of increased N supply by long residues at the stage of flag leaf emergence (GS37) which results in increased seed protein content, because N uptake by plants at GS37 is transferred to ear and seed development at GS59 to GS87 [75]. The association of the main stem elongation rate on day 42 after plant emergence with the final seed protein content was confirmed by the linear regression analysis which showed a small but significant influence. In addition, this correlation was further confirmed by Spearman correlation which showed a significantly negative correlation between final seed protein content and length of ears. Therefore, it seems that *long residues* contributed to higher main stem elongation rate on day 42 after plant emergence resulting in shorter ear length and higher seed protein content opposite to the *short residues*. Contrary, mulch diversity was not found to impact protein content. The effect of residue size on grains’ protein content should be further investigated as it is of great interest for both farmers and food processors.

In line with our hypothesis, it has already been shown that *long residues* had higher final C:N ratios, C, and N in comparison to *short residues*. In addition, it has been shown that there were no significant differences in residue dry mass loss or in crop yield between different residue size. Therefore, we reach the conclusion that *long residues* are characterized by an enhanced potential to provide nutrients to soil microbes and to the next crop at the end of the growing season than the *short residues*, without a yield penalty. At the same time, *long residues* provide better coverage of soil surface with organic material. Considering results were derived from a one growing season incubation experiment, it is highly possible the iteration of the practice of using *long residues* as mulch in successive crops could result in continuous enrichment of soil with nutrients, increase of soil organic matter, better physical conditions on soil surface, improvement of soil microbial community, and higher cost effectiveness in comparison to *short residues*. Further long-term research is needed to confirm this hypothesis.

The AMF root colonization was not significantly different between treatments. However, mean values were increased with increasing number of residue plant species. Regression analysis, considering the species richness (17, 12, 6, 1 and 0) of the different residue types and *Control*, showed a statistically significant influence of species richness in AMF root colonization, although the predictor could explain only a small variation of the response. Nevertheless, the fact that an increase in plant residue species richness on the soil surface is responsible for even a small increase in AMF root colonization of the crop plants is very important in terms of agricultural economic performance. This was consistent with previous finding by Burrows and Pfleger [22] who observed increasing AMF sporulation with increasing number of species of cover crop plants, which was attributed to increased number of AMF species triggered by the increased cover crop plant diversity. In our experiment the potential increase in AMF colonization in barley plant roots was triggered plainly by the deposition of cover crops cultivated elsewhere. This was an indication that residue mulch diversity may influence AMF root colonization and possibly the soil microbial community in general, which is in accordance with other researchers’ observations [20,21,76]. Nevertheless, it seems that the influence of residue diversity on soil microbial activity is greater at the early stage of decomposition due to functional complementarity in microbial communities which reduces competition in comparison to residues of lower diversity [19], but further investigation is needed.

### Conclusions

Dry mass loss of plant residue mulch was significantly affected by residue chemical composition. It was higher in residues of lower NDF and with lower N content. Moreover, the final quality of residues was highly affected by residue size, resulting in higher fertilization capacity of *long size* residues than of *short size* one. Soil K and Zn content were found to be significantly affected by residue quality and size, respectively, at later decomposition stages. Treatments of higher initial C:N ratio provided higher amounts of soil K, while *short residues* provided more Zn. Crop yield was not affected by residue quality or size. *Long residues* supported significantly higher rates of barley stem elongation than *short residues* at the stage of rapid stem elongation where N availability is determined for a high seed protein content. There were indications that mulches with *long residues* increased seed protein content, which is a key result important for both farmers and grain processors. The Arbuscular Mycorrhizal Fungi root colonization was higher but not significantly different in treatments with higher plant species richness, which indicates a possible effect of diverse mulch in soil microbial community. In summary, *long residue* mulches composed of diverse mixtures of plant species have enhanced residual fertilization capacity at the end of the growing season than short residues with no deleterious effect on crop yield. Further long-term research is needed to investigate the effect of continuous application of plant residue mulches on the enrichment of soil nutrient content, increase of soil organic matter, and improvement of soil microbial diversity.

## Acknowledgments

Authors would like to thank the contributors of the DiverseForages Project and Dr Anna Tomson in person who kindly provided plant residue material. Data supporting the results reported in this paper are openly available from the University of Reading Research Data Archive at https://doi.org/10.17864/1947.000395 [77].

## Availability of data and material

Not applicable.

## Code availability

Not applicable.

## Authors’ contributions

Conceptualization, D.G.; Experimental design, D.G., and M.L.; Experimental work, D.G.; Laboratory analyses, D.G.; Data analysis, D.G.; Data curation, D.G., M.L., and M.T.; Supervision, M.L., and M.T.; Writing – original draft, D.G.; Writing – review and editing, D.G., M.L., and M.T.

## Ethics approval

Not applicable.

