## Supplemental Information for "Fragment size and diversity of mulches affect their decomposition, nutrient dynamics, and soil microbiology"

**Appendices**

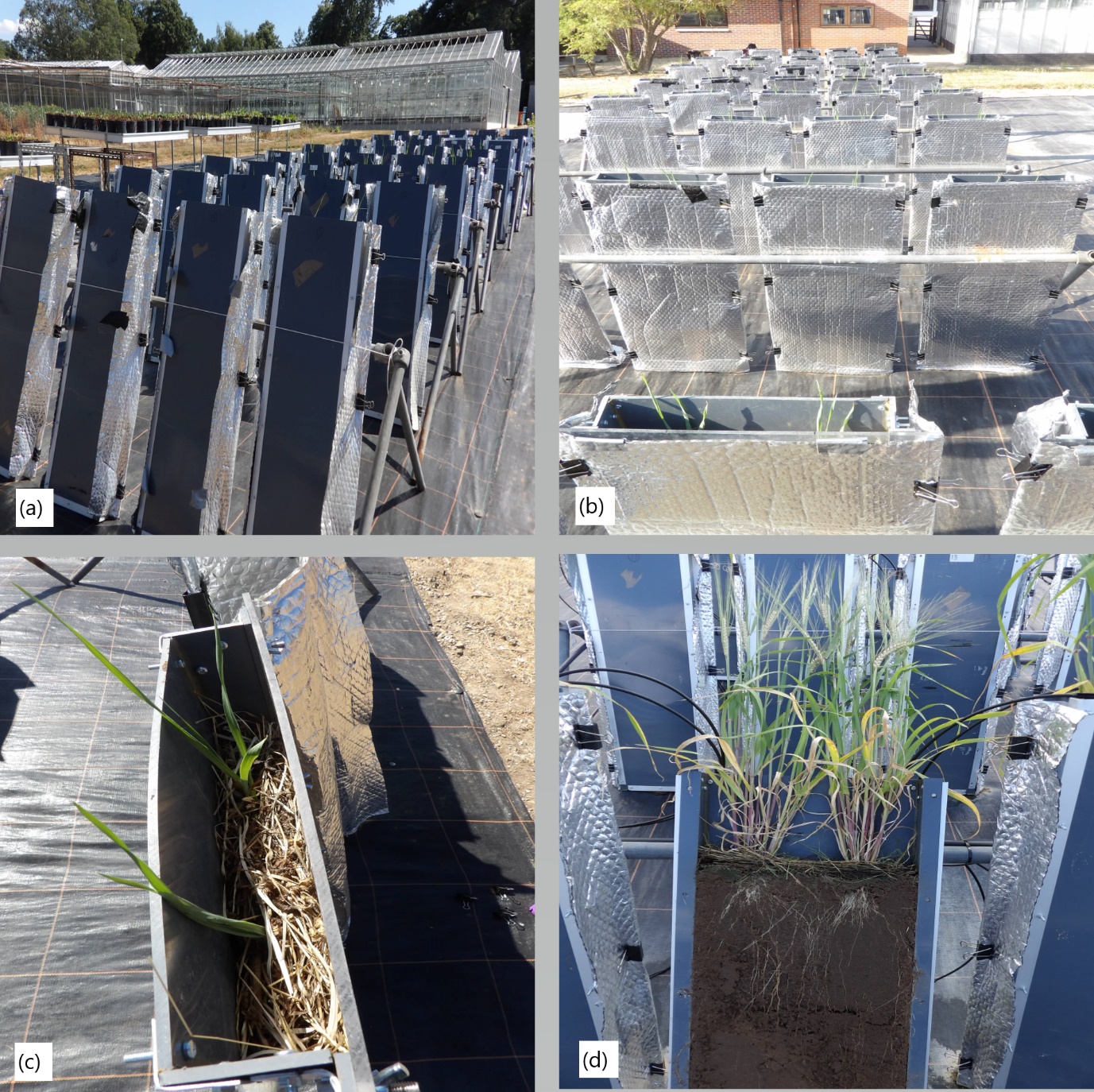

**Fig. S1** a) the back side of the rhizotrons settled at the outdoor, b) the front side covered with aluminum foil, c) a view from above with the growing plants, d) rhizotron with the glass removed to allow sampling.

**Table S1** Diverse forage mixture species selection list (P: Perennial Ryegrass; S: Smart Grass; B: Biomix; H: Herbal)

| **Species** | **Latin** | **P** | **S** | **B** | **H** |
| --- | --- | --- | --- | --- | --- |
| Perennial Ryegrass | *Lolium perenne* L. | ✓ | ✓ | ✓ | ✓ |
| Timothy | *Phleum pratense* L. |  | ✓ | ✓ | ✓ |
| Cocksfoot | *Dactylis glomerata* L. |  |  | ✓ | ✓ |
| Festulolium | - |  |  | ✓ | ✓ |
| Tall Fescue | *Festuca arundinacea* Schreb. |  |  |  | ✓ |
| Meadow Fescue | *Festuca pratensis* Huds. |  |  | ✓ | ✓ |
| Red Clover | *Trifolium pratense* L. |  | ✓ | ✓ | ✓ |
| White Clover | *Trifolium repens* L. |  | ✓ | ✓ | ✓ |
| Alsike Clover | *Trifolium hybridum* L. |  |  | ✓ | ✓ |
| Sweet Clover | *Melilotus* spp. |  |  |  | ✓ |
| Black Medick | *Medicago lupulina* L. |  |  | ✓ |  |
| Lucerne | *Medicago sativa* L. |  |  | ✓ |  |
| Sainfoin | *Onobrychis spp.* |  |  |  | ✓ |
| Birdsfoot Trefoil | *Lotus corniculatus* L. |  |  |  | ✓ |
| Plantain | *Plantago lanceolata* L. |  | ✓ | ✓ | ✓ |
| Chicory | *Cichorium intybus* L. |  | ✓ | ✓ | ✓ |
| Yarrow | *Achillea millefolium* L. |  |  |  | ✓ |
| Burnet | *Sanguisorba minor* Scop. |  |  |  | ✓ |
| Sheep’s Parsley | *Petroselenium crispum* Mill. |  |  |  | ✓ |

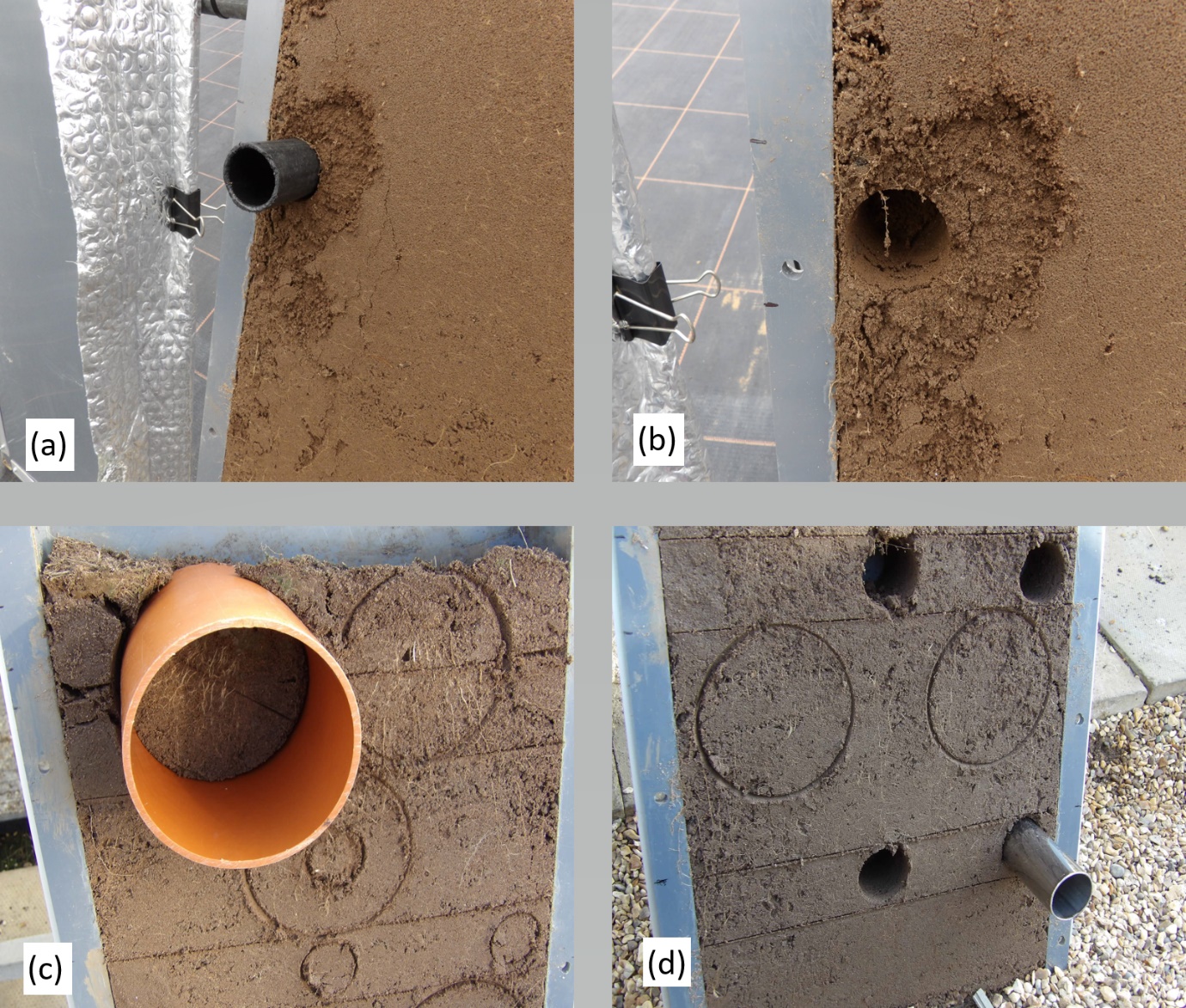

**Fig. S2** a) and b) 1^st^ sampling period (from the left side of the rhizotrons), c) and d) 2^nd^ sampling period.

**Table S2** Tukey’s post-hoc test for significant differences (p-value < 0.05) of initial C:N ratio, % C, and % N content of plant residues between different types. The residue types were: Perennial ryegrass *(****P****)* (1 plant species), Smart Grass *(****S****)* (6 species), Biomix *(****B****)* (12 species), and Herbal *(****H****)* (17 species)

| **C:N ratio** | | | **% C** | | | **% N** | | |
| --- | --- | --- | --- | --- | --- | --- | --- | --- |
| **Difference of Levels** | **T-Value** | **Adjusted P-Value** | **Difference of Levels** | **T-Value** | **Adjusted P-Value** | **Difference of Levels** | **T-Value** | **Adjusted P-Value** |
| *P - B* | -5.05 | <0.001 | *P - B* | 3.85 | 0.003 | *P - B* | 5.38 | <0.001 |
| *P - H* | -3.14 | 0.020 | *P - H* | 4.06 | 0.002 | *P - H* | 3.90 | 0.003 |
| *S - H* | 4.40 | 0.001 |  |  |  | *S - H* | -3.50 | 0.008 |
| *S - P* | 7.54 | <0.001 |  |  |  | *S - P* | -7.39 | <0.001 |

**Table S3** Initial C:N ratio, % C, and % N content for the different types of residues (N = 8). The residue types were: Perennial ryegrass *(****P****)* (1 plant species), Smart Grass *(****S****)* (6 species), Biomix *(****B****)* (12 species), and Herbal *(****H****)* (17 species)

| **Variables** | **C:N ratio** | | **% C** | | **% N** | |
| --- | --- | --- | --- | --- | --- | --- |
| **Types** | **Mean** | **StDev** | **Mean** | **StDev** | **Mean** | **StDev** |
| *B* | 30.420 | 3.640 | 41.229 | 0.256 | 1.373 | 0.178 |
| *H* | 28.069 | 1.801 | 41.207 | 0.195 | 1.473 | 0.091 |
| *P* | 24.199 | 2.268 | 41.644 | 0.160 | 1.734 | 0.170 |
| *S* | 33.502 | 1.650 | 41.400 | 0.237 | 1.238 | 0.055 |

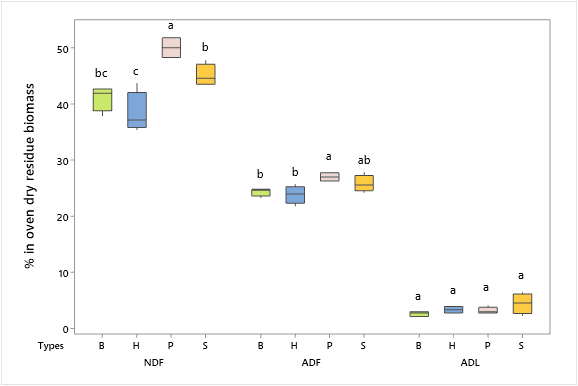

**Fig. S3** Box and whiskers plots of plant residue initial % NDF, % ADF, and % ADL for the different residue types. The residue types were: Perennial ryegrass *(****P****)* (1 plant species), Smart Grass *(****S****)* (6 species), Biomix *(****B****)* (12 species), and Herbal *(****H****)* (17 species). NDF = Neutral Detergent Fiber (Hemicellulose, Cellulose and Lignin), ADF = Acid Detergent Fiber (Cellulose and Lignin), and ADL = Acid Detergent Lignin (Lignin). Types that do not share a common letter are significantly different (p < 0.05)

**Table S4** Initial NDF, ADF, and ADL content (%) of the different residue types (N = 4). The residue types were: Perennial ryegrass *(****P****)* (1 plant species), Smart Grass *(****S****)* (6 species), Biomix *(****B****)* (12 species), and Herbal *(****H****)* (17 species). NDF = Neutral Detergent Fiber (Hemicellulose, Cellulose and Lignin), ADF = Acid Detergent Fiber (Cellulose and Lignin), and ADL = Acid Detergent Lignin (Lignin)

| **Variable** | **%NDF** | | **%ADF** | | **%ADL** | |
| --- | --- | --- | --- | --- | --- | --- |
| **Type** | **Mean** | **StDev** | **Mean** | **StDev** | **Mean** | **StDev** |
| *B* | 41.160 | 2.230 | 24.359 | 0.708 | 2.616 | 0.451 |
| *H* | 38.360 | 3.640 | 23.857 | 1.575 | 3.333 | 0.607 |
| *P* | 50.053 | 1.893 | 27.027 | 0.764 | 3.161 | 0.582 |
| *S* | 45.084 | 1.932 | 25.798 | 1.472 | 4.441 | 1.821 |

**Table S5** Tukey’s post-hoc test for significant differences (p-value < 0.05) of initial % NDF, ADF, hemicellulose, and cellulose content of residues between the different types of residues. The residue types were: Perennial ryegrass *(****P****)* (1 plant species), Smart Grass *(****S****)* (6 species), Biomix *(****B****)* (12 species), and Herbal *(****H****)* (17 species). NDF = Neutral Detergent Fiber (Hemicellulose, Cellulose and Lignin), ADF = Acid Detergent Fiber (Cellulose and Lignin), and ADL = Acid Detergent Lignin (Lignin)

| **% NDF** | | | | **% ADF** | | | |
| --- | --- | --- | --- | --- | --- | --- | --- |
| **Difference of Levels** | **Difference of Means** | **T-Value** | **Adjusted P-Value** | **Difference of Levels** | **Difference of Means** | **T-Value** | **Adjusted P-Value** |
| *P - B* | 8.90 | 4.98 | 0.002 | *P - B* | 2.66 | 3.15 | 0.036 |
| *P - H* | 11.69 | 6.55 | <0.001 | *P - H* | 3.17 | 3.75 | 0.013 |
| *S - H* | 6.72 | 3.76 | 0.012 |  |  |  |  |
| *S - P* | -4.97 | -2.78 | 0.069 |  |  |  |  |
| **% Hemicellulose** | | | | **% Cellulose** | | | |
| **Difference of Levels** | **Difference of Means** | **T-Value** | **Adjusted P-Value** | **Difference of Levels** | **Difference of Means** | **T-Value** | **Adjusted P-Value** |
| *P - B* | 6.77 | 5.98 | <0.001 | *P - H* | 3.34 | 2.57 | 0.099 |
| *S - B* | 4.31 | 3.81 | 0.012 |  |  |  |  |
| *P - H* | 8.35 | 7.37 | <0.001 |  |  |  |  |
| *S - H* | 5.89 | 5.20 | 0.001 |  |  |  |  |

**Table S6** Tukey’s post-hoc test for significant differences (p-value < 0.05) of final C:N ratio, % C, and % N content of residues between treatments (*HL, HS, BL, BS, SL, SS, PL, PS*). The residue types were: Perennial ryegrass *(****P****)* (1 plant species), Smart Grass *(****S****)* (6 species), Biomix *(****B****)* (12 species), and Herbal *(****H****)* (17 species). Residue treatments including four plant diversity mixtures (types) and two plant residue fibre sizes (short of 1.5 cm *(S),* and long of 30cm *(L)*)

| **C:N ratio** | | | **% C** | | | **% N** | | |
| --- | --- | --- | --- | --- | --- | --- | --- | --- |
| **Difference of Levels** | **T-Value** | **Adjusted P-Value** | **Difference of Levels** | **T-Value** | **Adjusted P-Value** | **Difference of Levels** | **T-Value** | **Adjusted P-Value** |
| *PL - BL* | 3.01 | 0.093 | HL - BS | 3.61 | 0.026 | *BS - BL* | -3.24 | 0.058 |
| *PL - BS* | 4.22 | 0.006 | PL - BS | 4.15 | 0.007 | *HS - BL* | -3.56 | 0.029 |
| *PS - BS* | 3.11 | 0.076 | SL - BS | 4.04 | 0.010 | *PS - BL* | -3.69 | 0.021 |
| *SL - BS* | 3.26 | 0.055 | HS - HL | -3.60 | 0.027 | *HL - BS* | 3.27 | 0.055 |
| *PL - HS* | 3.78 | 0.017 | PL - HS | 4.14 | 0.008 | *SL - BS* | 3.00 | 0.096 |
|  |  |  | SL - HS | 4.03 | 0.010 | *HS - HL* | -3.59 | 0.027 |
|  |  |  | PS - PL | -3.10 | 0.078 | *PS - HL* | -3.73 | 0.020 |
|  |  |  | SL - PS | 2.99 | 0.097 | *SL - HS* | 3.32 | 0.049 |
|  |  |  |  |  |  | *SL - PS* | 3.46 | 0.036 |

**Table S7** Final C:N ratio, % C, and % N of the different treatments (*HL, HS, BL, BS, SL, SS, PL, PS*) (N = 4). The residue types were: Perennial ryegrass *(****P****)* (1 plant species), Smart Grass *(****S****)* (6 species), Biomix *(****B****)* (12 species), and Herbal *(****H****)* (17 species). Residue treatments including four plant diversity mixtures (types) and two plant residue fibre sizes (short of 1.5 cm *(S),* and long of 30cm *(L)*)

| **Variables** | **C:N ratio** | |  | **% C** | | **% N** | |
| --- | --- | --- | --- | --- | --- | --- | --- |
| **Treatments** | **Mean** | **StDev** | **Initial – final C:N** | **Mean** | **StDev** | **Mean** | **StDev** |
| *BL* | 14,690 | 2,320 | 15.7 | 27.90 | 4.83 | 1.897 | 0.103 |
| *BS* | 13,079 | 0,651 | 17.3 | 17.02 | 3.65 | 1.303 | 0.279 |
| *HL* | 16,180 | 2,250 | 11.9 | 30.73 | 4.83 | 1.903 | 0.216 |
| *HS* | 13,666 | 0,281 | 14.4 | 17.07 | 5.16 | 1.244 | 0.356 |
| *PL* | 18,700 | 2,960 | 5.5 | 32.79 | 4.30 | 1.762 | 0.121 |
| *PS* | 17,221 | 1,824 | 7.0 | 21.01 | 4.28 | 1.218 | 0.193 |
| *SL* | 17,422 | 1,652 | 16.1 | 32.37 | 4.95 | 1.853 | 0.173 |
| *SS* | 16,150 | 1,607 | 17.3 | 22.65 | 9.10 | 1.385 | 0.442 |

**Table S8** Significant differences in soil available NH_4_^+^ content (mg/kg of oven dry soil) in samples taken on day 70 and on day 137 after mulch application, from 5-10 cm and from 50-55 cm depths, and two plant residue fibre sizes (short of 1.5 cm *(S),* and long of 30cm *(L)*) (N = 64)

| **Factors** | **Factor levels** | **Mean** | **StDev** | **F-value** | **P-value** | **Sign. diff.** |
| --- | --- | --- | --- | --- | --- | --- |
| Time | day70 | 1.302 | 0.356 | 46.65 | <0.001 | a*** |
|  | day137 | 1.678 | 0.379 |  |  | b*** |
| Depth | 10 | 1.689 | 0.280 | 52.13 | <0.001 | a*** |
|  | 55 | 1.292 | 0.428 |  |  | b*** |
| Time*Size | day70 *L* | 1.248 | 0.321 | 3.00 | 0.087 | a |
|  | day70 *S* | 1.357 | 0.385 |  |  | a |
|  | day137 *S* | 1.638 | 0.345 |  |  | b |
|  | day137 *L* | 1.719 | 0.412 |  |  | b |

*Combinations not shown in the table are not significant with all the others. Mean values that do not share a common letter are significantly different between them. Means with *** are very highly significant different, with ** are highly significant different, with * are significant different, with no * are nearly significant different*

**Table S9** Tukey’s post-hoc test for significant differences (p-value < 0.05) of soil NH_4_^+^ content for Size*Time interaction for *long* (*L*) and *short residues* (*S*), for day 70 and for day 137 after mulch application, from both 5-10 and 50-55 cm depths

| **Difference of Size*Time Levels** | **T-Value** | **Adjusted P-Value** | **Difference of Size*Time Levels** | **T-Value** | **Adjusted P-Value** |
| --- | --- | --- | --- | --- | --- |
| (*L* day137) - (*L* day70) | 6.05 | <0.001 | (*S* day70) - (*L* day137) | -4.65 | <0.001 |
| (*S* day137) - (L day70) | 5.01 | <0.001 | (*S* day137) - (*S* day70) | 3.61 | 0.003 |

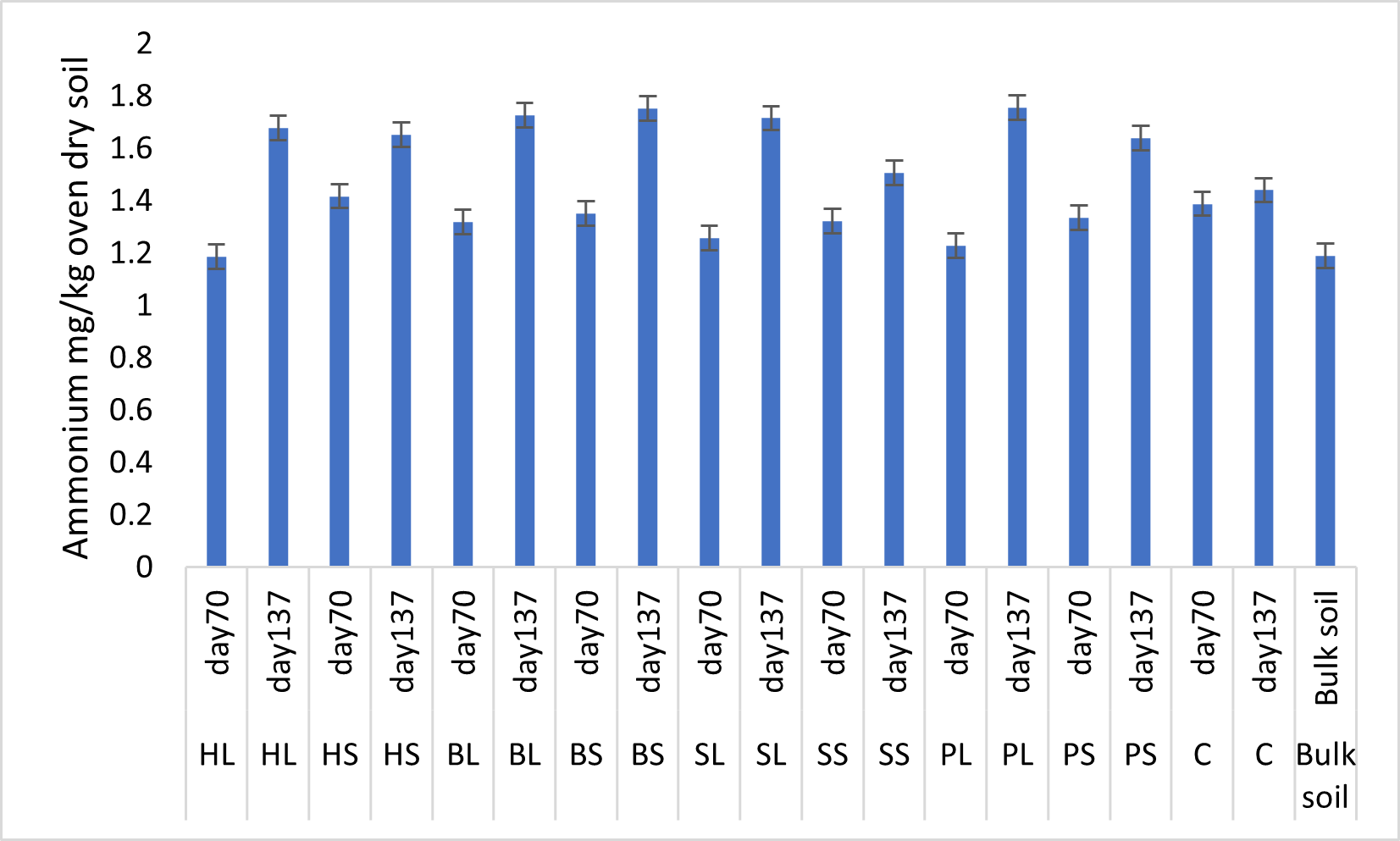

**Fig. S4** Soil NH_4_^+^ content (mg/kg of oven dry soil) from the different treatments (*HL, HS, BL, BS, SL, SS, PL, PS, and* *Control*) at both 5-10 and 50-55 cm depths, in September (day 70) and in November (day 137 after mulch application). The residue types were: Perennial ryegrass *(****P****)* (1 plant species), Smart Grass *(****S****)* (6 species), Biomix *(****B****)* (12 species), and Herbal *(****H****)* (17 species). Residue treatments including four plant diversity mixtures (types) and two plant residue fibre sizes (short of 1.5 cm *(S),* and long of 30cm *(L)*). Control treatment (*C*) was with no residues. Means and bars of one standard error from the mean are depicted. The bulk soil value is also quoted for comparisons

**Table S10** Soil available macronutrient (P, K, and Mg) contents (mg/kg of oven dry soil) in samples taken on day 70 and on day 137 after mulch application, from 0-5 cm and from 20-25 cm depths

| **Element** | **Factors** | **Factor levels** | **N** | **Mean** | **±SD** | **F-value** | **P-value** | **Sign. diff.** |
| --- | --- | --- | --- | --- | --- | --- | --- | --- |
| P | Time | day70 | 64 | 387.415 | 26.612 | 3800.71 | <0.001 | a*** |
|  |  | day137 | 64 | 210.224 | 162.204 |  |  | b*** |
|  | Depth | 25 | 64 | 388.441 | 26.590 | 3889.23 | <0.001 | a*** |
|  |  | 5 | 64 | 209.198 | 161.117 |  |  | b*** |
| K (fi) | Time | day70 | 64 | 103.163 | 37.343 | 28.40 | <0.001 | a*** |
|  |  | day137 | 64 | 79.907 | 21.985 |  |  | b*** |
|  | Depth | 5 | 64 | 97.985 | 42.640 | 8.74 | 0.004 | a** |
|  |  | 25 | 64 | 85.084 | 15.821 |  |  | b** |
| Mg (tr) | Time | day70 | 64 | 66.096 | 14.648 | 48.11 | <0.001 | a*** |
|  |  | day137 | 64 | 58.259 | 11.703 |  |  | b*** |
|  | Depth | 25 | 64 | 72.640 | 10.606 | 372.26 | <0.001 | a*** |
|  |  | 5 | 64 | 51.715 | 6.902 |  |  | b*** |

*Combinations not shown in the table are not significant with all the others. Mean values that do not share a common letter are* *significantly different between them. Means with *** are very highly significantly different, with ** are highly significantly different, with * are significantly different, with no * are nearly significantly different. Where tr = results obtained after transformation of data to log_10_ or to optimal or rounded λ, fi = results obtained without transformation of data and further investigation is needed because normality and/or equality of variances of data were not satisfied even after transformation*

**Table S11** Significant differences in soil available micronutrient (Fe, Mn, Zn, and Cu) contents (mg/kg of oven dry soil) in samples taken on day 70 and on day 137 after mulch application, from 0-5 cm and from 20-25 cm depths (N = 64)

| **Element** | **Factors** | **Factor levels** | **Mean** | **±SD** | **F-value** | **P-value** | **Sign. diff.** |
| --- | --- | --- | --- | --- | --- | --- | --- |
| Fe | Depth | 5 | 306.880 | 16.924 | 589.62 | <0.001 | a*** |
|  |  | 25 | 246.376 | 10.859 |  |  | b*** |
| Mn | Depth | 5 | 52.568 | 2.838 | 107.49 | <0.001 | a*** |
|  |  | 25 | 46.414 | 3.528 |  |  | b*** |
| Zn (tr) | Time | day70 | 26.301 | 15.841 | 46.91 | <0.001 | a*** |
|  |  | day137 | 20.167 | 13.065 |  |  | b*** |
|  | Depth | 5 | 26.807 | 14.200 | 66.21 | <0.001 | a*** |
|  |  | 25 | 19.661 | 14.590 |  |  | b*** |
| Cu (fi) | Time | day137 | 5.199 | 2.363 | 23.94 | <0.001 | a*** |
|  |  | day70 | 4.322 | 1.320 |  |  | b*** |
|  | Depth | 5 | 6.280 | 1.723 | 287.38 | <0.001 | a*** |
|  |  | 25 | 3.240 | 0.244 |  |  | b*** |

*Combinations not shown in the table are not significant with all the others. Mean values that do not share a common letter are* *significantly different between them. Means with *** are very highly significantly different, with ** are highly significantly different, with * are significantly different, with no * are nearly significantly different. Where tr = results obtained after transformation of data to log_10_ or to optimal or rounded λ, fi = results obtained without transformation of data and further investigation is needed because normality and/or* *equality of variances of data were not satisfied even after transformation*

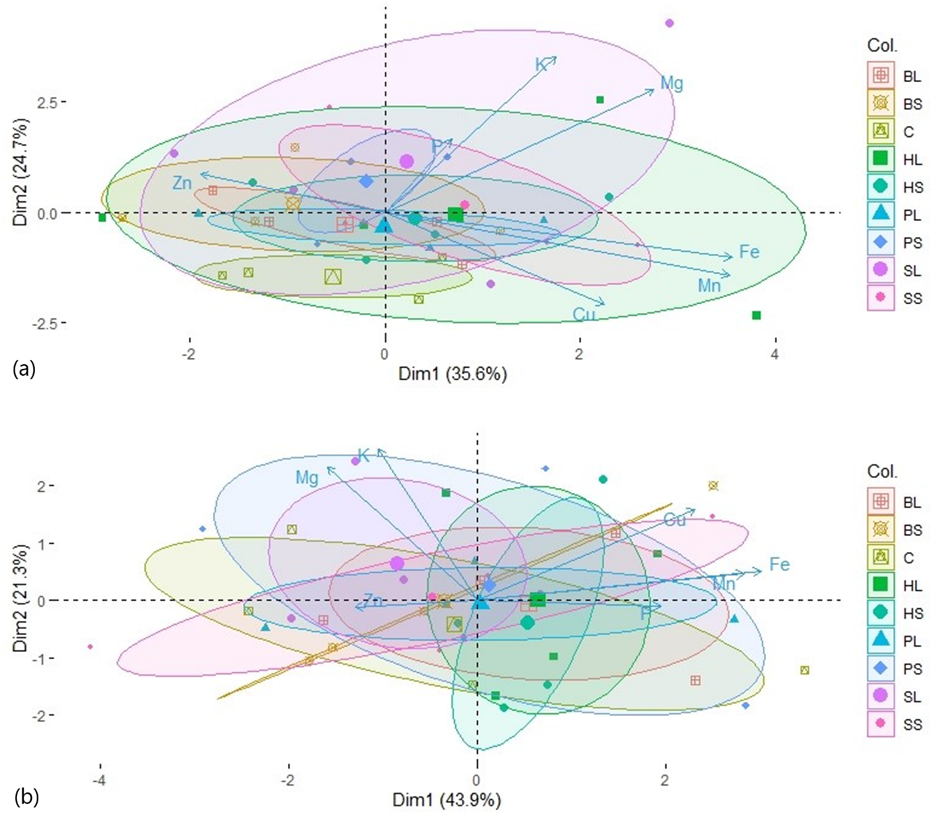

**Fig. S5** PCA ordination of nutrients P, K, Mg, Fe, Mn, Zn, and Cu as affected by the treatments *HL, HS, BL, BS, SL, SS, PL, PS,* and *Control (C)* on day 70 after mulch application (a) at 0-5 cm, and (b) at 20-25 cm depth. The residue types were: Perennial ryegrass *(****P****)* (1 plant species), Smart Grass *(****S****)* (6 species), Biomix *(****B****)* (12 species), and Herbal *(****H****)* (17 species). Residue treatments including four plant diversity mixtures (types) and two plant residue fibre sizes (*short* of 1.5 cm *(S),* and *long* of 30cm *(L)*). Control treatment (*C*) was with no residues. Large points depict mean values, while ellipses depict confidence intervals (ɑ = 0.05) of mean values for each treatment. The percentages represent data variation explained by the two first Principal Components (Dim1 and Dim2), bottom axis represents Dim1 normalized score, and left axis represents Dim2 normalized score

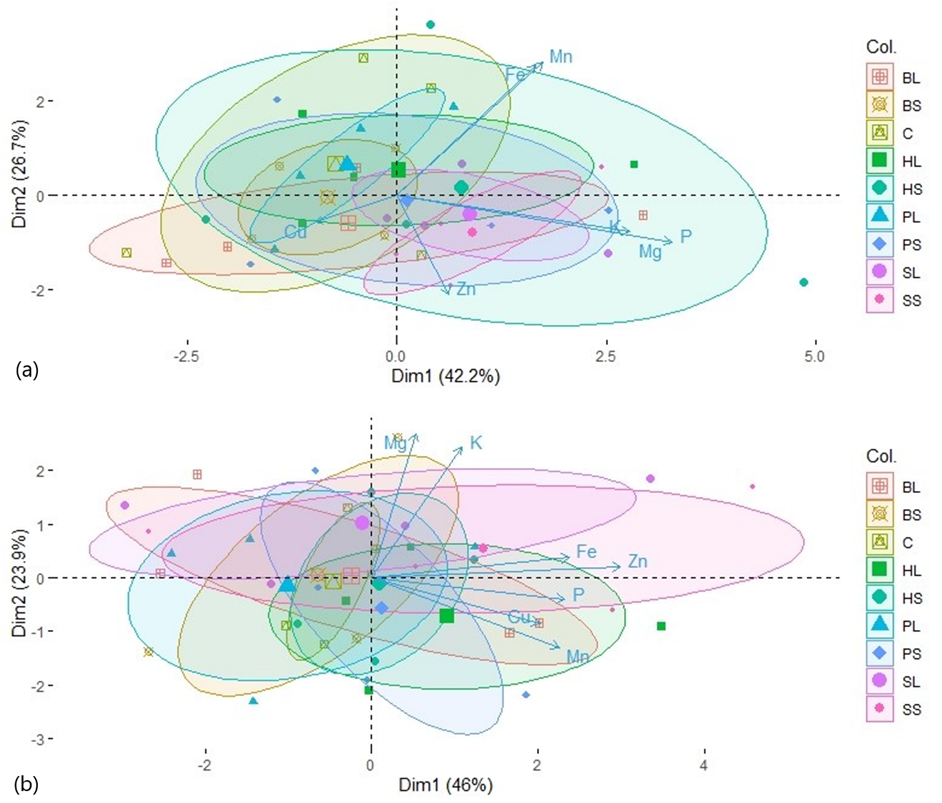

**Fig. S6** PCA ordination of nutrients P, K, Mg, Fe, Mn, Zn, and Cu as variables for the treatments *HL, HS, BL, BS, SL, SS, PL, PS,* and *Control* (*C*) concerning measurements (a) at 0-5 cm, and (b) at 20-25 cm depth, on day 137 after mulch application. The residue types were: Perennial ryegrass *(****P****)* (1 plant species), Smart Grass *(****S****)* (6 species), Biomix *(****B****)* (12 species), and Herbal *(****H****)* (17 species). Residue treatments including four plant diversity mixtures (types) and two plant residue fibre sizes (*short* of 1.5 cm *(S),* and *long* of 30cm *(L)*). Control treatment (*C*) was with no residues. Large points depict mean values, while ellipses depict confidence intervals (ɑ = 0.05) of mean values for each treatment. The percentages represent data variation explained by the two first Principal Components (Dim1 and Dim2), bottom axis represents Dim1 normalized score, and left axis represents Dim2 normalized score

**Table S12** Barley seed protein content of the different treatments (*BL, BS, C, HL, HS, PL, PS, SL, and SS*) (N = 4). The residue types were: Perennial ryegrass *(****P****)* (1 plant species), Smart Grass *(****S****)* (6 species), Biomix *(****B****)* (12 species), and Herbal *(****H****)* (17 species). Residue treatments including four plant diversity mixtures (types) and two plant residue fibre sizes (short of 1.5 cm *(S),* and long of 30cm *(L)*). Control treatment (*C*) was with no residues (N = 4)

| **Treatment** | **Mean** | **StDev** | **Treatment** | **Mean** | **StDev** |
| --- | --- | --- | --- | --- | --- |
| BL | 7.179 | 0.264 | PL | 6.930 | 0.234 |
| BS | 6.908 | 0.212 | PS | 7.087 | 0.691 |
| C | 6.995 | 0.545 | SL | 7.314 | 0.061 |
| HL | 7.877 | 1.271 | SS | 7.193 | 0.155 |
| HS | 7.126 | 0.480 |  |  |  |

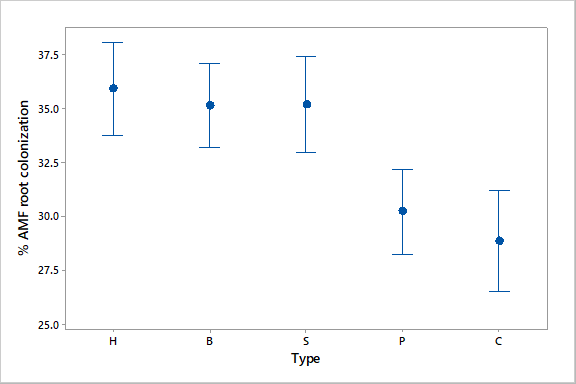

**Fig. S7** Interval plot of AMF root colonization (%) in all residue types and *Control* 137 days after mulch application (N = 8). The residue types were: Perennial ryegrass *(****P****)* (1 plant species), Smart Grass *(****S****)* (6 species), Biomix *(****B****)* (12 species), and Herbal *(****H****)* (17 species). Means and bars of one standard error from the mean are depicted

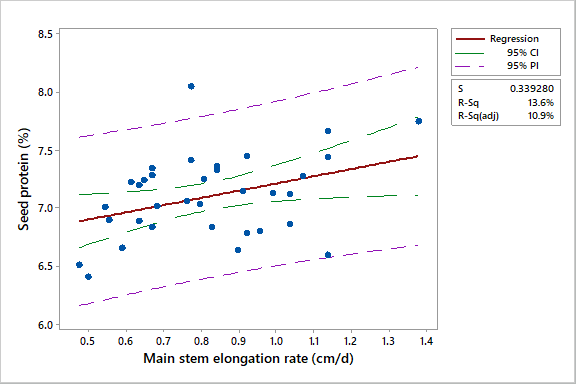

**Fig. S8** Fitted line plot of linear regression analysis of seed protein content (%) at harvest time with main stem elongation rate (MSER) (cm d^-1^) of barley plants on day 42 after plant emergence (red line). Confidence interval is enclosed by green dashed lines, and prediction interval by purple dashed lines. Confidence level = 95.0. Regression equation: Seed protein (%) = 6.589 + 0.6228 MSER
